# Uncovering the spatial landscape of molecular interactions within the tumor microenvironment through latent spaces

**DOI:** 10.1101/2022.06.02.490672

**Authors:** Atul Deshpande, Melanie Loth, Dimitrios N. Sidiropoulos, Shuming Zhang, Long Yuan, Alexander Bell, Qingfeng Zhu, Won Jin Ho, Cesar Santa-Maria, Daniele Gilkes, Stephen R. Williams, Cedric R. Uytingco, Jennifer Chew, Andrej Hartnett, Zachary W. Bent, Alexander V. Favorov, Aleksander S. Popel, Mark Yarchoan, Lei Zheng, Elizabeth M. Jaffee, Robert Anders, Ludmila Danilova, Genevieve Stein-O’Brien, Luciane T. Kagohara, Elana J. Fertig

## Abstract

Recent advances in spatial transcriptomics (ST) enable gene expression measurements from a tissue sample while retaining its spatial context. This technology enables unprecedented *in situ* resolution of the regulatory pathways that underlie the heterogeneity in the tumor and its microenvironment (TME). The direct characterization of cellular co-localization with spatial technologies facilities quantification of the molecular changes resulting from direct cell-cell interaction, as occurs in tumor-immune interactions. We present SpaceMarkers, a novel bioinformatics algorithm to infer molecular changes from cell-cell interaction from latent space analysis of ST data. We apply this approach to infer molecular changes from tumor-immune interactions in Visium spatial transcriptomics data of metastasis, invasive and precursor lesions, and immunotherapy treatment. Further transfer learning in matched scRNA-seq data enabled further quantification of the specific cell types in which SpaceMarkers are enriched. Altogether, SpaceMarkers can identify the location and context-specific molecular interactions within the TME from ST data.

## 1 Introduction

The tumor microenvironment (TME) is the tissue region created and controlled by a tumor in its surroundings and plays a key role in tumorigenesis and therapeutic response in cancer [1–4]. The TME contains tumor cells, stroma, blood vessels, and immune cells as well as cells from the resident tissue [4]. A thorough understanding of the molecular profile of individual cells and the impact of inter-cellular interactions in the TME is crucial for distinguishing the determinants of tumor progression [5–7] and precision medicine strategies[3, 8–11].

Advances in single-cell technologies have led to the development of spatially resolved transcriptomics (ST) which captures the transcriptome *in situ* [12] and thus allows us to study the spatial relationship between the various cell populations within the TME as well as their relationship with the tumor cells. For example, the 10X Visium spatial transcriptomic technology allows us to resolve tissue heterogeneity at near single-cell resolution (from one to ten cells per spot). The technique has been applied to characterize the cellular and molecular composition of tumors [13–15]. Robust analysis pipelines for cell-based analysis and cellular deconvolution have been proposed to model the cellular composition of spatial-transcriptomics data [16–20] and cellular phenotypes within each spot [21]. While spot deconvolution methods can infer linear combinations of molecular markers that are reflective of cellular co-localization, new computational methods are needed to characterize the molecular changes resulting from cell-cell interaction at a genome-wide scale.

Many analysis pipelines for Visium ST rely on latent space methods for cellular deconvolution to overcome the mixture of cells at each spot. In this paper, we present the SpaceMarkers algorithm which leverages spatial overlapping latent features to infer interactions between cell types or biological processes represented by latent patterns. We demonstrate the use of our method using latent features inferred from CoGAPS, a Bayesian non-negative matrix factorization approach. We selected CoGAPS based on its robustness for single-cell RNA-seq data [22, 23]. We apply CoGAPS followed by SpaceMarkers to distinct tumor samples to benchmark its performance on both immune alterations resulting from tumor infiltration in the lymph node and on neoplastic cell alterations resulting from immune infiltration of intra-tumoral lesions. Briefly, we first demonstrate that the CoGAPS approach can delineate the established cellular architecture and associated molecular changes to those cells resulting from a pancreatic ductal adenocarcinoma (PDAC) that has metastasized to the lymph node. We then demonstrate how multi-resolution factorization of Visium data can further uncover the molecular pathways of distinct lesions within breast cancer and enable the inference of intra-tumor heterogeneity of immune cell interactions using SpaceMarkers. Finally, integration of Visium data with scRNA-seq data of a liver sample from a patient treated with immunotherapy allows us to determine the specific cell subtypes in which the identified SpaceMarkers genes are enriched. Altogether, our extension to latent space analysis enables us to simultaneously infer cellular architecture and model molecular changes resulting from spatially interacting biological processes.

## 2 Results

### 2.1 Interactions between overlapping latent features delineate inter-cellular interactions in ST data

Here we present SpaceMarkers, a novel bioinformatics algorithm for identifying genes associated with cell-cell interactions in ST data. SpaceMarkers is an extension of latent space analysis that leverages spatially overlapping latent features associated with distinct cellular signatures to infer the genes associated with their interaction (Figure 1). Fundamentally, this inference relies on estimation of spatially resolved linear latent features representative of cellular signatures in the ST data. That is, the latent feature information is characterized by continuous weights corresponding to each spatial coordinate in the ST data. We denote these continuous weights as the patterns in the ST data. The inputs to the SpaceMarkers algorithm are the ST data matrix and spatially resolved patterns learned through latent space analysis, and the output is a list of genes associated with the interaction between each pair of spatially overlapping patterns. The first stage of the algorithm involves the identification of each pattern’s region of influence and subsequently the region of pattern interaction (Figure 1A.; see also Methods). If a pattern has a nonzero value at a point, we hypothesize that its influence extends to its neighboring region but rapidly decreases with increasing distance. We model this by spatially smoothing the patterns using a Gaussian-kernel based approach (see Methods). Subsequently, we identify outlier values of smoothed patterns by testing it against a null-distribution obtained by identical smoothing of spatially permuted pattern values. We denote the region corresponding to these outlier values as the region of influence of the pattern. Furthermore, two patterns are deemed to be interacting in the region with overlapping influence from both patterns. We hypothesize that genes associated with the spatially overlapping influence from two patterns represent changes in molecular pathways due to the interaction between the biological features of the associated cells. Therefore, we devise the second stage of the SpaceMarkers algorithm to rank genes exhibiting higher activity levels in the interaction region relative to regions with exclusive influence from each pattern (Figure 1B.; see also Methods). To this end, we perform a non-parametric statistical test followed by posthoc analysis to identify these genes which constitute the SpaceMarkers output.

**Figure 1.**
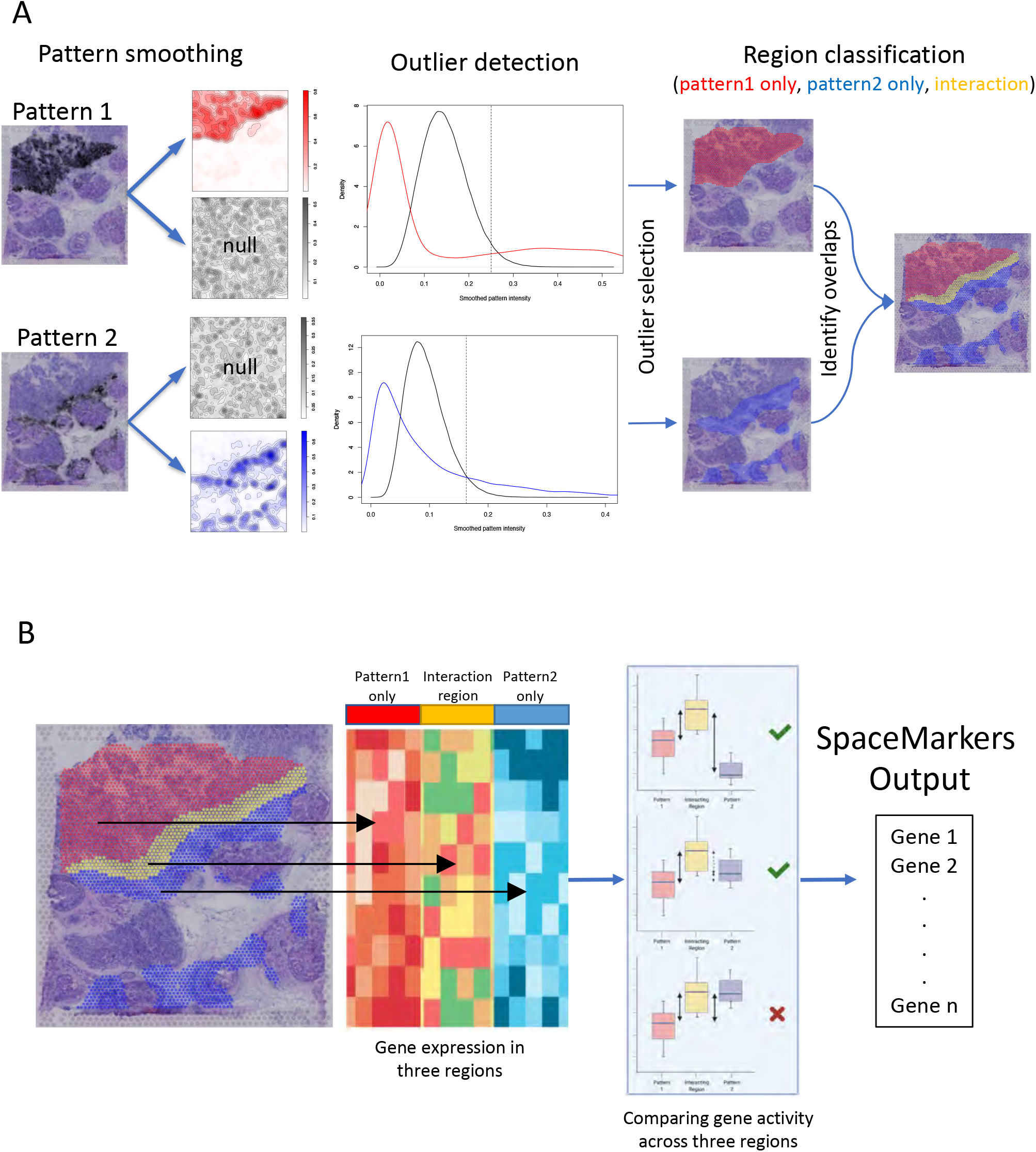
SpaceMarkers identifies genes associated with cell-cell interaction using spatially overlapping patterns. **A.** Identifying interaction region: The input to the SpaceMarkers algorithm are spatially resolved latent features resulting from latent space analyses (e.g. CoGAPS patterns). The images on the left show the intensity levels of two spatially resolved CoGAPS patterns. For each pattern, the SpaceMarkers algorithm first identifies regions of influence (red and blue spots, respectively) using a Gaussian-kernel based outlier detection method. The patterns are deemed to be interacting in the region with overlapping influence (yellow spots) from both patterns. It also identifies regions with mutually exclusive influence from each pattern (red and blue spots). **B.** Identifying SpaceMarkers genes: The second stage of the SpaceMarkers algorithm performs a non-parametric Kruskal-Wallis statistical test with posthoc analysis on the gene expression data in the three regions (pattern 1 only, pattern 2 only, and interaction region) to identify molecular changes due to cell-cell interaction. The output is a list of genes associated with the pattern interaction (see Methods).

In the examples demonstrated here, the spatial data is obtained using the spot-based 10x Visium spatial transcriptomics technology [12] with 1-10 cells per spot. SpaceMarkers is readily applicable to spot-based ST data with regions of influence and interaction defined as sets of spots in which one or two patterns respectively have influence as identified by the Gaussian-kernel based approach. We use CoGAPS Bayesian nonnegative matrix factorization [24, 25] for identifying the latent features associated with cellular signatures. When two patterns have overlapping influence in the same region of the tissue, we assume an interaction between these patterns in this interaction region. We provide a differential expression (DE) mode for SpaceMarkers to quantify genes with enhanced expression in a region with overlapping influence from two patterns when compared to regions with exclusive influence from individual patterns. Further we extend this approach to provide a “residual” mode — which identifies genes that have significantly higher residual error between the original ST data and its estimated fit from the CoGAPS model in the region with overlapping influence from two patterns when compared to the regions with exclusive influence from each pattern. We hypothesize that the residual mode detects the nonlinear effects of intercellular interaction more precisely by accounting for the underlying linear latent features to mitigate confounding effects from variations in the cell population density and cell types with common markers. Thus, the SpaceMarkers algorithm infers both simple molecular changes in the “DE” mode as well as more precise nonlinear molecular changes in the “residual” mode in regions with overlapping influence from patterns associated with different cell signatures. We denote such patterns with concurrent influence in a region as “spatially interacting” patterns. The reliance on latent space patterns from CoGAPS enables the further ability to integrate SpaceMarkers learned from ST data in corresponding single-cell data using transfer learning from projectR [23, 26] to refine the specific cells in which these molecular changes occur. While the method is developed for the example of latent space patterns in ST data from CoGAPS to define cellular signatures, it is generally applicable to patterns from any of a number of linear latent feature factorization approaches available in the literature.

### 2.2 SpaceMarkers identifies molecular changes from tumor immune interactions associated with metastatic pancreatic cancer cells invading the lymph node

In the first example, we applied SpaceMarkers on Visium ST data from a pancreatic cancer metastasis to the lymph node in a patient who received neoadjuvant GVAX vaccination (see Figure 2). More specifically, this sample is characterized by the presence of metastatic PDAC, immune cell aggregates, and germinal centers of B-cell maturation (Figure 2A.). Analysis of the H&E imaging from the lymph node region used to generate the ST data identifies a region of the tissue in which the metastatic PDAC intersects the immune cells surrounding the germinal center. On factorizing this data using CoGAPS we obtain ten latent patterns based only on the expression data (Fig. S1, Methods). By matching pattern activity levels learned from the data with the independent histological annotations, we observe that CoGAPS can distinguish metastatic PDAC in Pattern 6 from immune cells in the surrounding lymph node tissue in Pattern 9 (Fig. 2B.).

**Figure 2.**
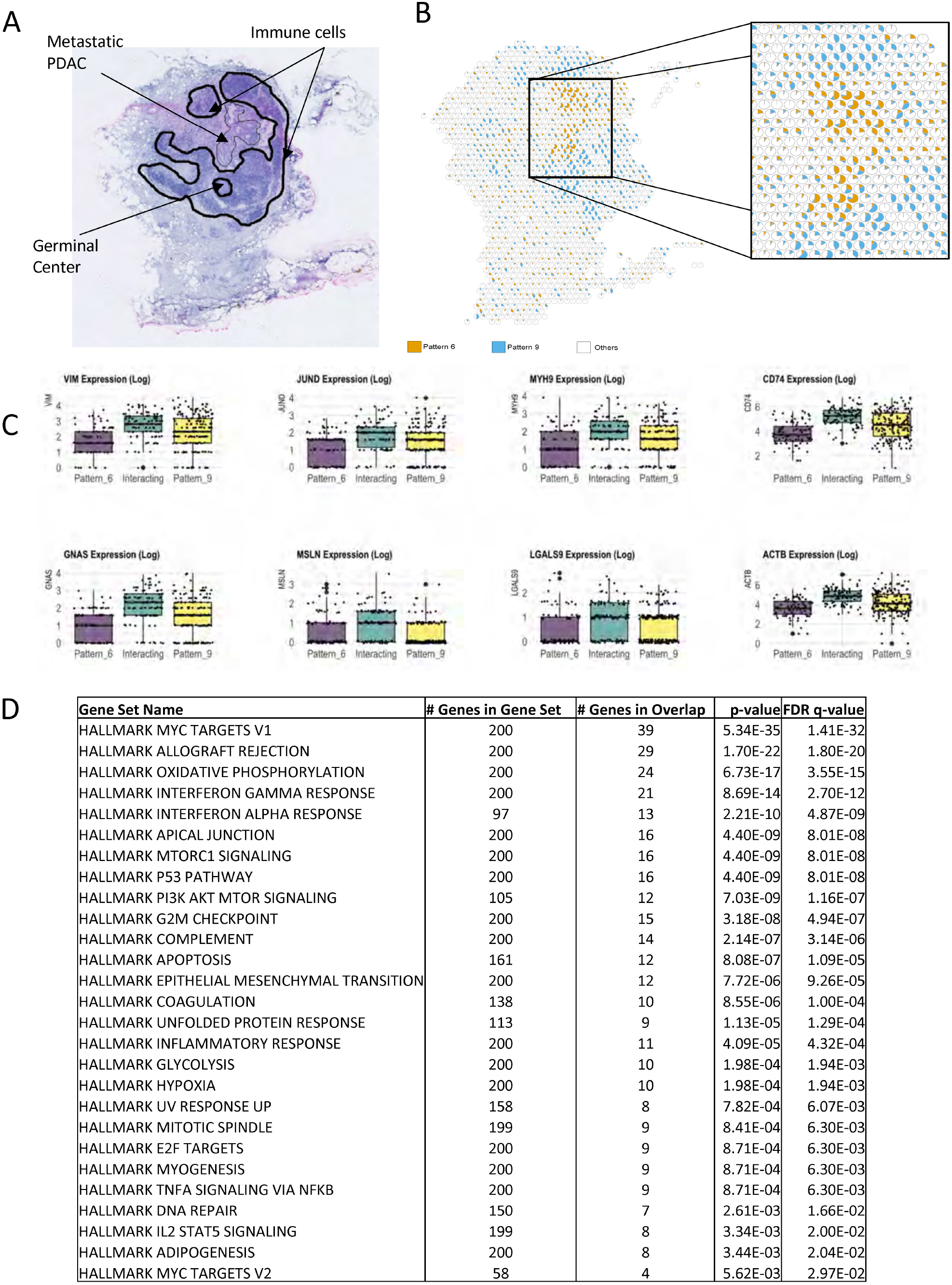
SpaceMarkers identifies molecular changes associated with immune-metastatic pancreatic cancer interaction in the lymph node **A.** H&E staining of a peritumoral pancreatic lymph node with metastasis from PDAC (arrow) and annotated germinal center and immune cells (dark lines). **B.** Visualization of the relative activity in the CoGAPS patterns associated with metastatic PDAC (Pattern 6) and immune cells in the lymph node (Pattern 9). Each spot is represented as a pie chart with fractional gene expression at the location aggregated over the all genes for Pattern 6 (red), Pattern 9 (blue) and the other patterns (white). **C.** Boxplots of the expression of selected genes showing higher expression levels in the interaction region of Pattern 6 and Pattern 9 compared to the regions with exclusive influence from Pattern 6 and Pattern 9 respectively. **D.** Table showing Hallmark gene set pathways significantly overrepresented in the region of interaction between Pattern 6 and Pattern 9, with size of overlap and FDR value (see Table S1 for KEGG and Biocarta pathways).

We further analyzed the spatial activity of the metastatic PDAC (Pattern 6) and immune (Pattern 9) patterns to identify regions of overlapping influence to associate with metastasis-immune interaction. We represent the spatial variation in the activity levels of Pattern 6 and Pattern 9 in relation to all the other patterns in each spot (Fig. 2B.). This proportional analysis of patterns enables us to observe a spatial overlap between the regions where Pattern 6 and Pattern 9 are active. However, we hypothesize that a pattern has influence in a spot even with zero pattern activity but high pattern activity levels in the neighboring spots. We deem this overlapping region as the interaction region between the two patterns. We then use the SpaceMarkers algorithm to define the gene expression changes that occur from metastasis-immune interaction in the spots where the two patterns have overlapping influence. Due to the limited number of spots where the two patterns have overlapping influence, we define SpaceMarkers based upon differential expression. This analysis identifies 1442 genes which exhibit higher average expression in the interaction region with overlapping influence from the two patterns compared to spots where only metastatic PDAC in Pattern 6 or immune cells in Pattern 9 have exclusive influence (see Methods for details of the statistical test, Table S2 for complete gene list with the associated statistics).

Figure S1 shows the expression heatmap of the SpaceMarkers genes in spots belonging to regions with exclusive influence from the metastatic PDAC Pattern 6, exclusive influence from the immune cell Pattern 9, and overlapping influence from both patterns in metastasis-immune interaction. In all cases, the interactions are associated with changes in extracellular matrix genes, including notably genes associated with cytoskeleton regulation (*TMSB10, TMSB4X, CFL1, MARCKSL1*), the myosin pathway (*MYL6, MYH9, MYL12B*), actin regulation (*ACTB, ACTN4, CAPG, LCP1, SPTBN1*), the matrix metallopeptidase family (*MMP9, MMP12*), galectin genes (*LGALS1, LGALS4, LGALS9, LGALS3BP*), collagen (*COL1A2, COL3A1, COL4A1, COL4A2, COL18A1, COL6A2*), and cell adhesion (*MSLN, ITGB4, ITGB6, ADRM1*). The SpaceMarkers include genes reflecting cell death in the increased expression of ribosomal protein genes associated with immune response through expression of HLA family genes, immunogoblulins, interleukins, cytokines, chemokines, the interferon pathway *IFITM2*, and immune function. This immune response is counterbalanced by changes to pathways associated with enhanced invasion in cancer cells, including *JUNB, JUND, VIM*.

To further elucidate the molecular pathways associated with the metastasis-immune interaction in the lymph node, we performed gene set overrepresentation analysis (Figure 2D., Table S1) from the Hallmark, KEGG, and Biocarta molecular pathways using the Molecular Signatures Database (MSigDB) [27–29]. As seen in Fig. 2D., Hallmark pathways related to allograft rejection, interferon gamma, and interferon alpha are all overrepresented in the pathway analysis for the SpaceMarkers genes, and hence in the region of overlap between the immune and metastatic PDAC patterns. This confirms activation of the immune response for tumor rejection at the interface between the metastatic PDAC and the immune cells in the lymph node observed at the gene level. Likewise, we observe overrepresentation in the epithelial to mesenchymal signaling and pathways consistent with the invasive process in the metastatic PDAC cells, further supported by the enrichment of the apical junction consistent with the changes to the extracellular matrix suggested by the gene-level SpaceMarkers analysis.

The DE mode of SpaceMarkers is applicable when the available latent features provide only a partial reconstruction of the original ST data matrix. However, the differential expression of a marker in the interaction region could be due to nonlinear effects because of two intercellular interactions or due to confounders such as variable cell populations in each spot and different co-localized cell types having common markers. In the examples to follow, we mitigate these confounding effects by using the residual error between the raw expression and its reconstruction from the CoGAPS patterns, which capture the effect of both variations in cell population density as well as variations in individual marker expression.

### 2.3 SpaceMarkers identifies markers of tumor-immune interactions in invasive breast ductal carcinoma through residual space analysis

While providing a means to detect molecular changes from cellular interactions in limited interaction regions, using differential expression statistics for SpaceMarkers could confound nonlinear effects from cell-cell interactions with expression changes resulting from increased density of co-localized cell types with shared gene markers. In cases where the interaction region extends across a greater number of spots, these confounding effects can be mitigated by using the residual error between the raw expression and its estimated fit from the CoGAPS model for the SpaceMarkers. This estimated fit will capture the effect of both variations in cell population density as well as variations in individual marker expression to refine the estimates of the nonlinear effects from cell-cell interactions. We apply this approach to identify the molecular pathways associated with tumor cell and immune interactions in ST data from a breast cancer sample that contains multiple ductal carcinoma in situ (DCIS) lesions, an invasive carcinoma lesion, immune cells, and stroma (Figure 3A.).

**Figure 3.**
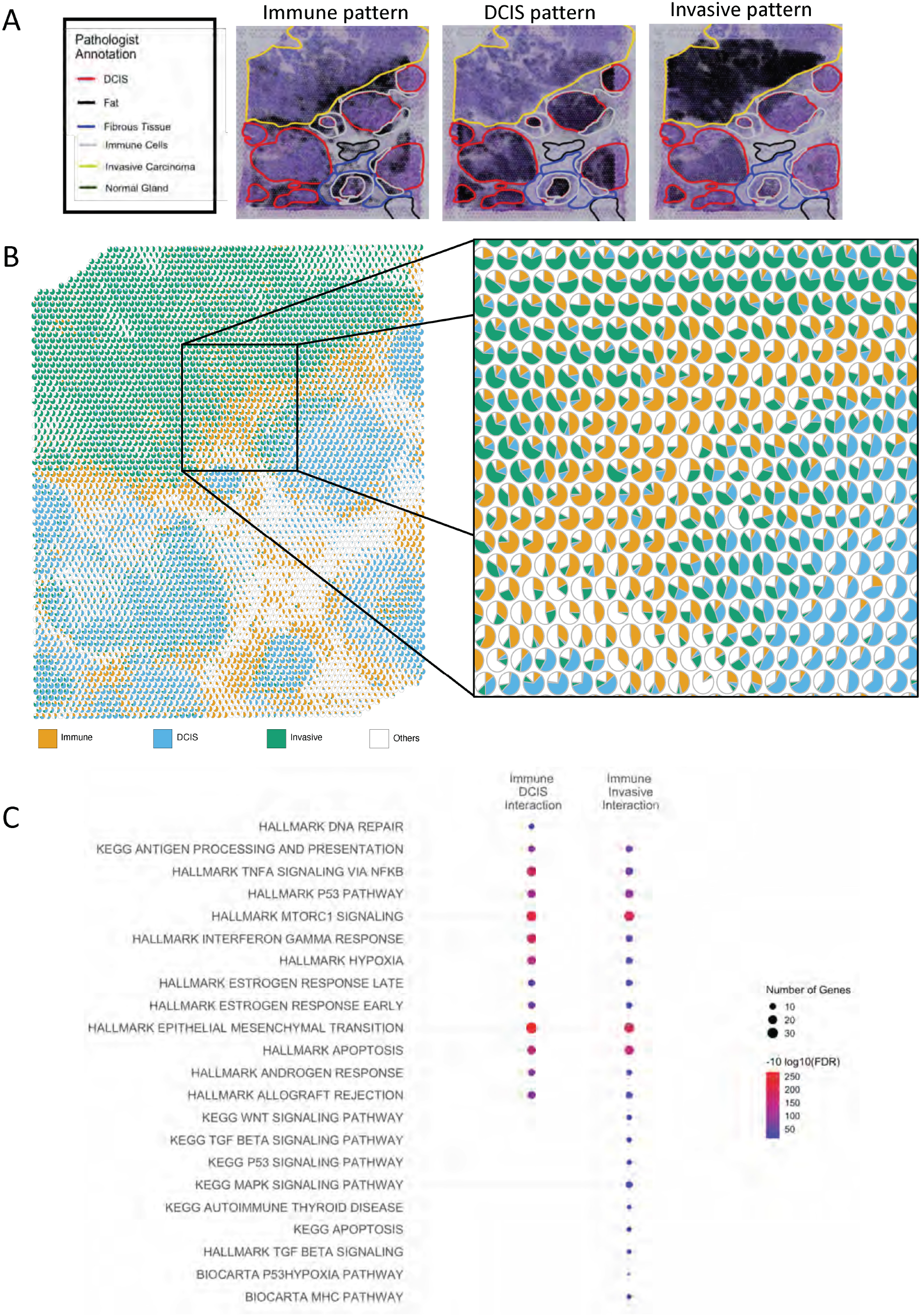
Low-resolution CoGAPS and SpaceMarkers analysis identifies markers of interaction between broad patterns in breast cancer tissue **A.** Images of the breast cancer tissue showing activity levels of the immune, DCIS, and invasive carcinoma patterns respectively overlaid on annotated H&E slides showing regions with invasive carcinoma, DCIS lesions, immune cells and stroma. **B.** Scatterpie visualization shows the relative activity levels and overlap between the invasive carcinoma, immune, and DCIS patterns. **C.** Enriched pathways associated with DCIS-immune interactions and cancer-immune interactions.

The visualization in Figure 3B. shows widespread spatial regions of interactions between immune and tumor cells at the boundaries of both the invasive carcinoma and the DCIS lesions. Whereas the pancreatic cancer sample in Figure 2 covered a smaller area with fewer spots (< 300) having tumor and immune influence respectively, we identify much larger regions (> 1000 spots) of influence from the immune, invasive carcinoma and DCIS cells (Figure 3B.). This larger number of spots enables us to estimate SpaceMarkers from CoGAPS residuals to distinguish the molecular changes in the invasive carcinoma from the DCIS lesions. Similar to our analysis of the metastatic pancreatic cancer data, we obtain latent features of the ST data from this breast sample using CoGAPS factorization. These latent features reveal histological annotations of invasive carcinoma, DCIS lesions, immune, and stromal regions estimated from the H&E stain (Figure 3A.).

Computing SpaceMarkers based upon the CoGAPS residuals identifies 461 genes associated with interaction between the immune and invasive carcinoma patterns and 413 markers of immune and DCIS pattern interaction (Supplemental File 2A), compared to up to 3736 immune-invasive carcinoma and 3036 immune-DCIS genes identified from applying a similar analysis based upon differential expression for the same FDR value (Supplemental File 2B). This reduction in the number of markers through the analysis of CoGAPS residuals relative to inference of SpaceMarkers through differential expression analysis is consistent with the isolation of specific nonlinear changes resulting from interactions between the cellular processes measured in the CoGAPS patterns using this mode. We note that 85 of the SpaceMarkers were associated with immune cell interactions in both the invasive carcinoma and DCIS regions. To further determine the molecular pathways activated through immune and tumor cell interactions in both regions, we performed gene set overrepresentation analysis from the Hallmark, Kegg, and Biocarta molecular pathways using the Molecular Signatures Database (MSigDB), with a selection of the pathways presented in Figure 3C. (see Table S3 for the complete list of pathways). We find that while certain pathways were enriched in both interactions (e.g., antigen processing and presentation, p53 pathway, Tnf-alpha signaling, mTorc1 signaling, epithelial to mesenchymal transition, Interferon Gamma response, hypoxia, and estrogen response early/late), others were enriched exclusively in Immune-DCIS (DNA repair) and Immune-Invasive (WNT signaling, MapK signaling, and TGF beta signaling) respectively. Note that a pathway enriched in both Immune-DCIS and Immune-Invasive Carcinoma interactions may have distinct gene subsets associated with each interaction. For example, it is readily evident that the Hallmark interferon gamma response gene set has a greater overlap with the SpaceMarkers of the Immune-DCIS interaction compared to the Immune-Invasive interaction.

### 2.4 Using SpaceMarkers with high-resolution CoGAPS reveals greater heterogeneity in intercellular interactions within the TME

In all cases presented, the SpaceMarkers inferred fundamentally depend on the resolution of the cellular processes inferred in the CoGAPS latent space analysis. Indeed, nonlinear interactions in interacting regions at a low resolution analysis may be further refined by increasing the dimensionality of the factorization on the ST data consistent with recent advances to multi-resolution matrix factorization [30]. We further performed a higher resolution CoGAPS analysis of the breast cancer data to test if the interaction region between two patterns and the associated SpaceMarkers genes are identified by increasing the dimensionality of the latent space analysis. In this higher dimensional analysis, CoGAPS identifies 16 distinct patterns associated with the diverse biological processes in the TME. The activity levels of a selection of the patterns overlaid on an H&E stained slide of the sample are shown in Figure 4A. (also see Figure S2). Although the higher number of patterns reveal greater heterogeneity of the biological processes in the sample by further resolving patterns identified in the low resolution analysis, it does not identify patterns specific to the interactions identified between the lower dimension patterns.

Although we do not associate each Visium spot with solely one pattern, studying the most dominant pattern in spots informs us of the dominant biological process at that location in the tissue as inferred by CoGAPS. Consequently, the same spots are associated with broader biological processes at the lower resolution and with more specific processes at a higher resolution. The alluvial plot in Figure 4B. shows the relationship between the most dominant low resolution and high resolution patterns at each spot.

**Figure 4.**
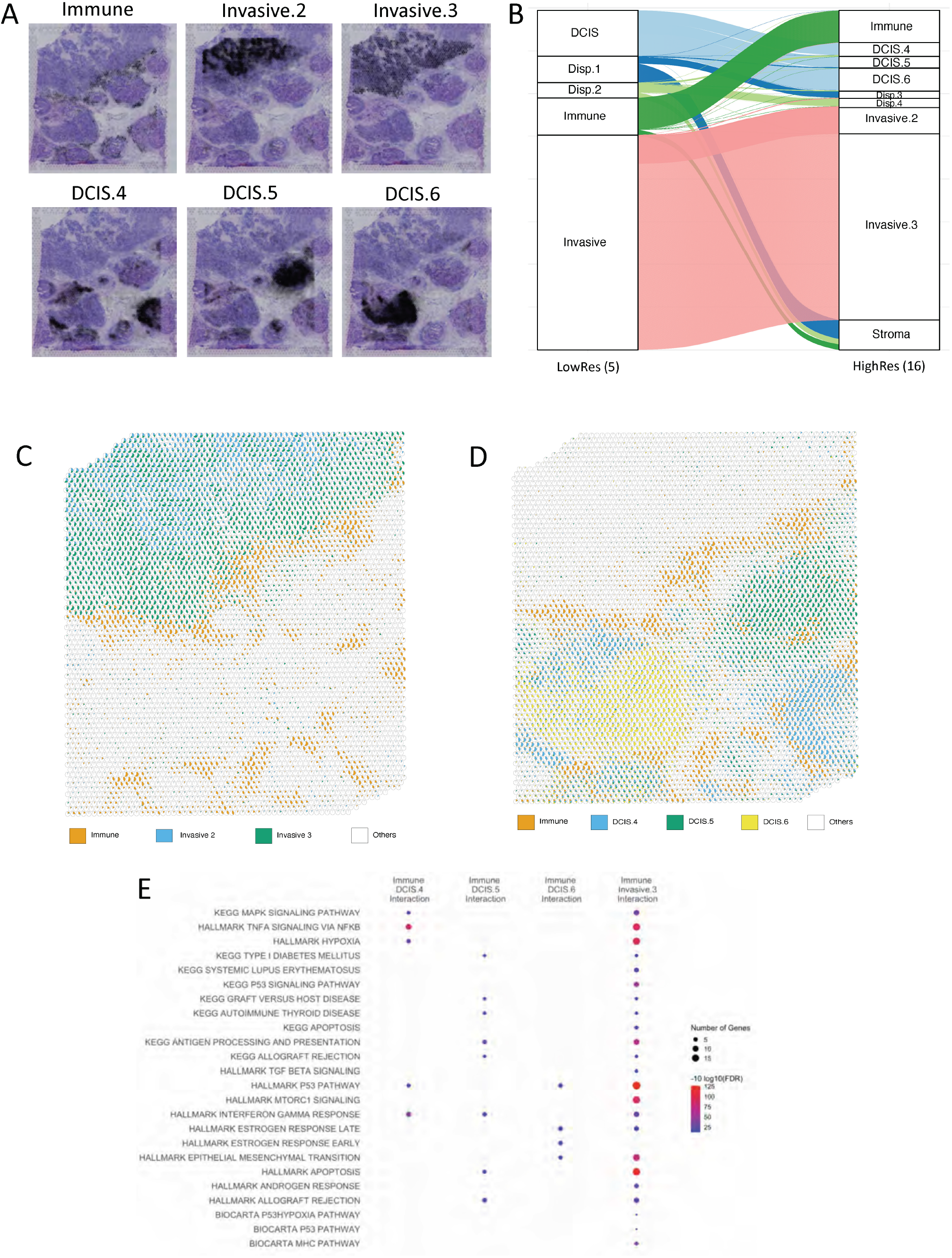
High-resolution CoGAPS and SpaceMarkers analysis of breast cancer tissue reveal greater heterogeneity in intercellular interactions. **A.** Multiple patterns associated with invasive carcinoma and DCIS regions identified in higher-resolution CoGAPS analysis with 16 patterns highlights the heterogeneity in the tumor and TME by further resolving the underlying pathology (see Figure S2 for remaining patterns). **B.** Alluvial plot showing the most dominant pattern associated with each spot using low-resolution and high-resolution CoGAPS respectively. Spots dominated by low resolution DCIS pattern are dominated by three distinct DCIS-related patterns associated with different lesions in the high-resolution analysis. Invasive pattern in low resolution resolves into three invasive carcinoma related patterns associated with varying levels of immune infiltration in the high-resolution analysis. **C.** Relative activity levels of immune patterns with two invasive patterns reveals that the immune and Invasive.2 patterns have no overlap, hence do not interact. Immune interaction with Invasive Carcinoma is captured through the overlap between Immune and Invasive.3 pattern. **D.** Relative activity levels of immune patterns with three DCIS patterns associated with separate lesions reveals distinct overlapping regions associated with each interaction. **E.** SpaceMarkers of Immune-DCIS and immune-Invasive interactions reveal functional heterogeneity of the enriched pathways mirroring the spatial heterogeneity revealed in B-E. (See Table S4 for complete list of gene sets)

For example, the single DCIS-related pattern in Figure 3A. resolves into multiple DCIS patterns, some of which are associated with individual DCIS lesions. Even within the single invasive carcinoma lesion, the low resolution invasive carcinoma pattern resolves into two distinct patterns, one of which is isolated to the interior of the invasive carcinoma and one which spans to the tumor-immune boundary. While the DCIS lesions and invasive carcinoma have universally high *ERBB2* and *ESR1* expression, evaluating the genes associated with the distinct patterns identifies heterogeneity in growth factor signaling pathways with enhanced *IGFBP3* expression in the DCIS.5 pattern, *FGFR4* expression in the DCIS.6 pattern, and *FGFR1* expression in the Invasive.2 carcinoma pattern (Figure S2, Table S5) We also see spots previously associated with the immune pattern or with dispersed patterns at the low resolution now being associated with a dominant pattern which can be associated with the stromal region. To further compare the enhanced resolution intra-tumor heterogeneity to tumor-immune interactions in the high resolution factorization, Figure 4C. shows relative pattern weights and overlap between the immune pattern and the two invasive patterns. It is clear that only one of the invasive patterns overlaps with the immune pattern, thus contributing to the tumor-immune interaction. Still, both of these interacting patterns contain a substantial numbers of spots that are isolated to the immune and invasive carcinoma region, respectively, suggesting that increasing the resolution of the factorization does not compensate for the estimation of nonlinear effects through the interaction statistic. Similarly, Figure 4D. shows relative pattern weights and overlap between the immune pattern and the three DCIS patterns. It logically follows that the overlapping regions of the distinct DCIS patterns are also distinct, and hence correspond to different molecular alterations from DCIS-immune interactions that will impact subsequent outgrowth of these distinct lesions.

For these interactions involving the immune pattern, we identify SpaceMarkers genes associated with the inter-pattern interactions as the genes having higher CoGAPS residuals in the interaction region compared to regions with exclusive influence from the individual patterns. Upon identification of statistically significant (*FDR* < 0.05) signaling pathways (see Supplementary Table S4) pertaining to interaction of the immune pattern with invasive carcinoma and DCIS patterns in the high-dimensional CoGAPS results and comparing them to those found in 5 dimensions, we find pathways common to all interactions and unique to specific pattern interactions. For example, we find 59 signaling pathways enriched due to immune-invasive carcinoma interaction in 5 dimensions as well as 16 dimensions. These include but are not limited to pathways related to epithelial-mesenchymal transition, apoptosis, antigen processing and presentation, hypoxia, p53 signaling, interferon alpha and gamma responses, and lastly targets of the oncogene MYC. However, the higher resolution analysis also reveals unique pathways relevant to specific immune-invasive carcinoma pattern interactions. We found pathways related to the cancer-immune interactions including those related to IL-5 and IL6 signaling, KRAS signaling, Toll-like receptor signaling and the CDC25 pathway exclusively when the dominant invasive carcinoma pattern (Invasive.3) interacts with the immune cells. Similarly, the distinct Immune-DCIS interactions reveal a heterogeneity in the enriched pathways which were not evident with a single DCIS pattern using low-resolution CoGAPS. Among the immune interactions with different DCIS lesions, the MapK signaling, Tnf alpha signaling, and hypoxia pathways, known to be mechanisms of resistance to endocrine and immunotherapies, are enriched exclusively in the Immune-DCIS.4 interaction, antigen processing, allograft rejection and autoimmunity related pathways are enriched exclusively in Immune-DCIS.5, and EMT pathway, and estrogen response early/late are exclusively enriched in the Immune DCIS.6 interaction. These pathways are consistent with the heterogeneity of subsequent outgrowth of these DCIS lesions, with successful activation of pathways associated with immune attack in DCIS.5 relative to the invasive processes observed in both DCIS.4 and DCIS.6.

### 2.5 Integrated ST and single-cell RNA-seq analysis identifies cell type specific molecular changes from immunotherapy treatment in hepatocellular carcinoma

In the examples so far, the SpaceMarkers statistic revealed molecular changes associated with intercellular interactions. Since SpaceMarkers relies on spot-based colocalization, it limits the ability to identify the cell subtypes in which these molecular changes were induced. Transfer learning allows us project new data into learned latent spaces, subsequently associating samples from the new data with known biology. We first factorize the ST data collected from a resected hepatocellular carcinoma (HCC) tumor after administration of a neoadjuvant cabozantinib and nivolumab therapy to obtain 9 CoGAPS patterns. Figure 5A. shows the individual tumor and immune associated patterns overlaid on an H&E stained image of the HCC tumor sample. As in the other examples, these tumor and immune patterns are spatially overlapping (Figure 5B.), and are deemed to be interacting in regions where they have overlapping influence. This analysis identifies two distinct tumor cell patterns, one of which spans all malignant regions in the sample (Pattern 2) and the other isolated to a specific region (Pattern 1) that has less co-localization of the immune cells (Pattern 8). The interaction between the immune cells and each of the tumor patterns learned through SpaceMarkers identifies enhanced expression of hepatocyte markers (*KRT18, SERPIN* family genes, *APOC2, CD24*), immune markers (*CD63, HLA* genes), and cell death markers (TNF pathway associated genes, ribosomal genes, *ANXA2*) consistent with killing of tumor cells through immune cells in the interaction between Patterns 2 and 8 (see Supplemental Table S5). In contrast, SpaceMarkers genes of the interaction between Patterns 1 and 8 identify fibroblast markers (Collagen coding genes, *MYL9*, *TAGLN*) consistent with a lack of successful immune attack and infiltration in this portion of the tumor.

**Figure 5.**
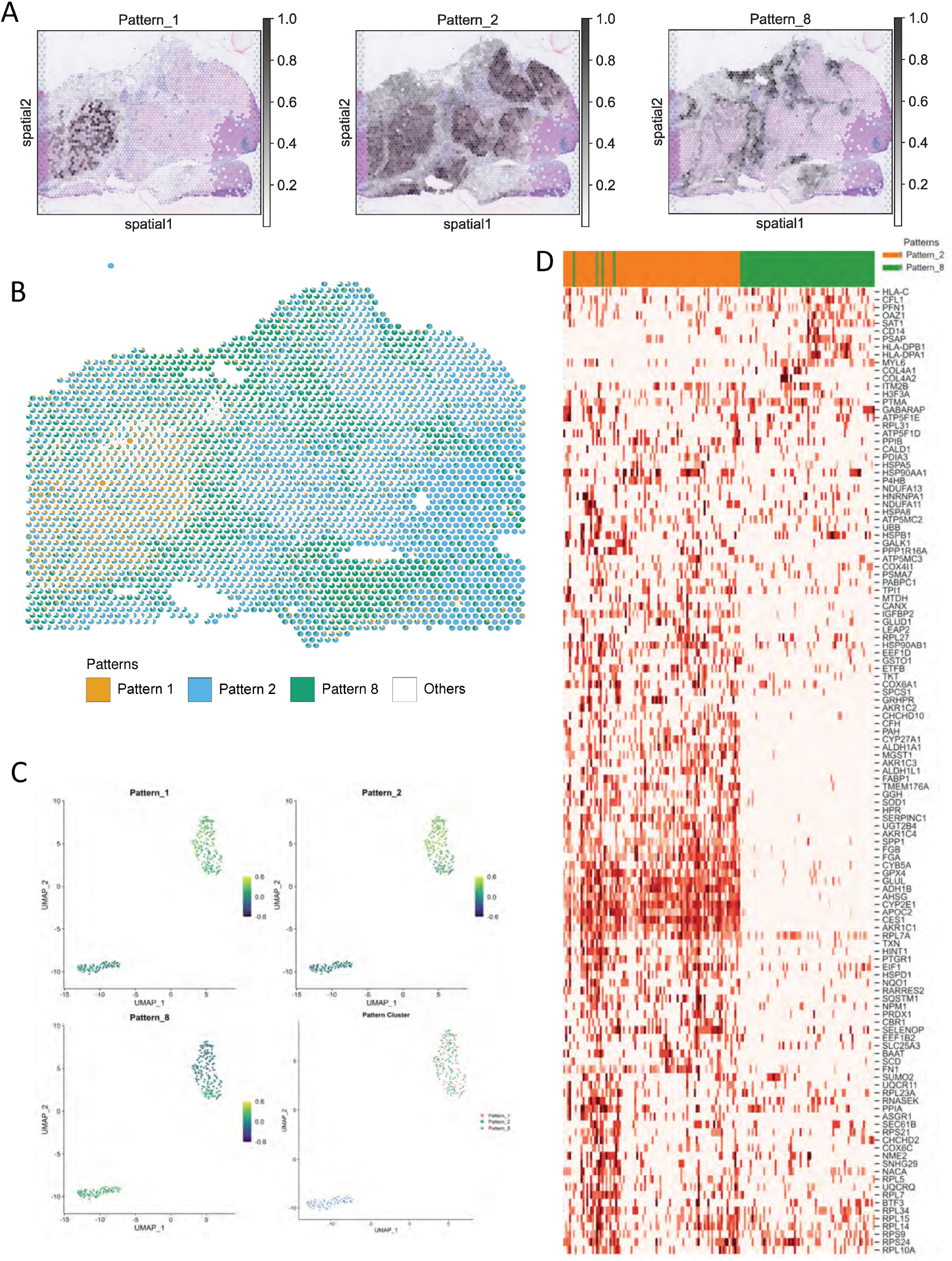
Contextualizing scRNAseq data using SpaceMarkers and transfer learning from matched ST-scRNAseq data in HCC (see also Figures S3 and S4). **A.** CoGAPS factorization reveals spatial patterns associated with tumor annotations of tumor and immune cells. (see Figure S3). **B.** Scatterpie visualization shows the relative pattern activity levels associated with the spatially overlapping tumor (orange) and immune (blue) patterns in each Visium spot using a pie chart (white represents activity from all other patterns). SpaceMarkers are genes exhibiting nonlinear effects in the residual space of the CoGAPS patterns in the region with tumor-immune overlap. **C.** Transfer learning of Patterns 1, 2, and 8 from ST data to matched scRNAseq data. Scatter plot shows projections of the spatial patterns onto individual cells in the scRNAseq data. Individual cells in the scRNAseq data are associated with the pattern having the highest projection in the cell. **D.** Expression heatmap of SpaceMarkers in tumor and immune cells from matched single-cell data from the same tumor provide the spatial context of the individual cells.

While the SpaceMarkers analysis of ST data suggests molecular changes associated with cell-cell interactions, this analysis alone does not pinpoint the precise cells in which these molecular changes occur. By transfer learning [23, 26] of these latent features into matched single-cell RNA-seq data from the same tumor, we can associate individual cells with specific patterns corresponding to tumor and immune signatures (Figure 5C.). This association can both identify whether a SpaceMarkers gene’s expression changes in tumor or immune cells, and also whether we can also predict the precise subpopulations of tumor and immune cells involved in intercellular interactions by observing the gene expression changes of the relevant SpaceMarkers in individual cells. From Figure 5D., we observe that changes in the expression of genes *SERPINC1, APOC2* and *ADH1B*, are induced in a subset of the cancer cells attributed to Pattern 2, whereas expression changes in gene *PFN1* and *CD14* are induced in a subset of the immune cells. A further subset of both Pattern 2 tumor cells and immune cells co-express *HSP90AA1* and ribosomal genes. Based on these gene expression patterns of the respective SpaceMarkers, we hypothesize that these individual cells are sourced from the tumor-immune boundary.

## 3 Discussion

We demonstrate how co-localization of multiple cellular processes within a single spot from ST data from Visium can be leveraged as an asset to infer molecular changes resulting from inter-cellular interactions. Specifically, this inference is enabled through SpaceMarkers, a novel algorithm for identifying genes associated with a spatial overlap between pairs of latent features that each represent distinct cellular processes. We accomplish this by first identifying a region of influence for each latent feature of interest. The two features are deemed to be interacting in spots where they have concurrent influence. While the focus in this paper is explicitly on cellular co-localization in spot-based technologies, we note that SpaceMarkers is readily applicable to alternative imaging-based ST technologies that achieve single-cell resolutions.

The SpaceMarkers algorithm can estimate molecular changes from spatially overlapping cellular processes in two ways — a default residual mode and a differential expression (DE) mode. We demonstrate that the DE mode is able to identify genes with significantly higher expression in the region where two latent features overlap. However, the DE mode is subject to confounding factors such as variable cell populations and marker association with multiple cell types. We mitigate these confounding effects in the residual mode, where we identify genes with significantly higher residual error between the original data and its reconstruction in the region of overlap between two latent features. However, this statistic requires a greater number of spots for robust analysis than the DE method. While we found that this requirement limited application of the residual model in the case with the smaller lymph node sample with PDAC metastasis, it was generally applicable to the other tumor-immune interactions in our sample cohort. Moreover, increased spatial resolution of the ST characterization or multi-omics methods for inferring cellular boundaries [31, 32] will enable broader application of this methodology for inter-cellular interactions. We note in the current algorithm if a cell type is entirely occurring within the interaction region, its marker genes will be inferred through SpaceMarkers. While not a direct molecular change in the input cell states, this co-localization exclusively in the interaction region may be a biological effect induced through the TME state induced by the inter-cellular interactions. Regardless, this effect is mitigated in the residual mode if some of the learned patterns are associated with that cell type.

We note that our inference of interactions between cellular processes is performed directly from latent space analyses of the ST data, without the need for additional reference datasets for single-cell resolution [16] or direct estimates of cellular deconvolution [18]. While our approach is generally applicable to linear latent space estimation methods, the results of our algorithm fundamentally depend on the latent space method selected for analysis of the ST data. We demonstrate the application of SpaceMarkers to 10x Visium ST data from different cancers and we identify markers associated with the interaction between latent features associated with different biological processes. In all cases, we observe that the Bayesian matrix factorization method CoGAPS [23, 24, 33] can learn latent features that distinguish regions with tumor and immune cells directly from the ST data without reliance on prior knowledge of marker genes, histology annotations, or spatial coordinates. Moreover, creating higher resolution CoGAPS analysis by increasing the number of latent features inferred from the ST data is able to further resolve the biological signatures, revealing the tissue heterogeneity. These higher dimensional patterns are independent of the interaction regions between the latent features inferred with SpaceMarkers at a lower dimension. This observation suggests that our approach indeed isolates non-linear effects due to inter-cellular interactions rather than unresolved latent features associated with specific cellular processes.

Due to our focus on tumor-immune cell interactions in our biological analyses, the current version of the SpaceMarkers algorithm admits only two overlapping latent features as input. However, this approach is generally applicable to inter-cellular inference from ST data across biological contexts. In many cases, multiple latent features are co-localized at the same spot. This could result in the same genes being associated with multiple interaction types, although we did not observe such effects in our case studies. Furthermore, many critical intercellular interactions such as cancer-associated fibroblast (CAF)-driven immunosuppression [34] result from possible colocalization of multiple cell phenotypes. To address this, future work should extend the application of SpaceMarkers to identify genes associated with multiple overlapping latent features. The use of SpaceMarkers on spot-based co-localization limits direct inference of the specific cell subtypes in which interactions induce molecular alterations. We demonstrate that transfer learning [23, 26] of the latent features inferred from CoGAPS analysis of the ST data into matched single-cell RNA-seq data enables us to define the precise cellular subpopulations with gene expression changes in each SpaceMarkers gene. Other approaches mitigate the need for paired data by coordinated expression changes between annotated pairs of ligands and receptors in both spatial and non-spatial single-cell data. While these approaches directly model the signaling process, they rely on the correspondence between gene expression and protein function and databases of ligand-receptor pairs [35]. Coupling spatial data with newer single-cell technologies that isolate interacting cells [36] can further enhance this inference.

Ultimately, the results of SpaceMarkers depends on the patterns inferred from the latent space method. Biological robustness of the SpaceMarkers statistic relies on the use of patterns associated with significant activity levels as well as a spatial overlap with other patterns of interest. For example, we analyzed the interaction of immune cells with one invasive carcinoma pattern out of the three invasive carcinoma patterns learned using high resolution CoGAPS analysis. We did not analyze the other two patterns because one was isolated away from the immune pattern and hence had no interactions, and although the other pattern had a spatial overlap with the immune pattern, it had much lower activity levels. For the residual mode to be effectively used, it is important not just to resolve the ST data into biologically meaningful latent features, but also to provide a good fit between the original ST data and its reconstruction from the latent features. In the absence of a good fit, the residual errors contain not just the nonlinear effects attributable to inter-feature interaction and the measurement error, but also the estimation errors resulting from an overly constrained factorization. In such cases, we recommend using the SpaceMarkers in the DE mode. Similarly, the utility of SpaceMarkers is diminished if the learned latent features do not correspond to individual cell phenotypes, or if markers of essential cell types are not represented by any of the learned latent features. Future work can overcome this limitation through with semi-supervised learning methods that use cell-type marker expression as a proxy for the latent feature input in the DE mode for SpaceMarkers.

When genes associated with cell-surface interactions and cytokine secretions are grouped together in a latent feature, the assignment of a single kernel-width parameter to the latent feature in the SpaceMarkers algorithm is inconsistent with the varying distances associated with these two types of intercellular interactions. Identification of intercellular interactions in such scenarios requires a mathematical framework for spatially resolved causal inference which models distinct cell types, varying ranges and gradients of influence for cytokine-secretions and surface interactions, and spatially resolved expression of individual genes. One such example is MESSI [37], which uses mixture-of-experts and multi-task learning approaches to predict the gene expression in a particular cell type with the help of signaling genes in neighboring cells. Future work integrating these methods with latent features in place of individual genes will both reduce the computational complexity and enhance the biological interpretability of these spatially aware network inference methods.

## 4 Methods

### 4.1 Sample collection, preparation, and storage

#### Invasive breast ductal carcinoma

The fresh frozen invasive breast ductal carcinoma was collected in 2011 and obtained from BioIVT. The tumor was stage IIA, ER Positive, PR Negative, Hercep Test 2+. The RNA quality of the sample, as measured with Bioanalyzer (Agilent) was RIN = 9.26. The sample was embedded in optimal cutting temperature (OCT) compound and immediately frozen. Cryosections of 10 *μ*m were placed on Visium Gene Expression slides (10x Genomics).

#### PDAC metastatic lymph node

The PDAC peritumoral lypmh node was surgically resected during curative surgery at the Johns Hopkins University. The lymph node was embedded in OCT and immediately frozen. Pathological examination of an H&E stained cryosection identified a PDAC metastasis to the lymph node. A cryosection of 10 *μ*m were placed on a Visium Gene Expression slide (10x Genomics)

#### HCC sample

The HCC sample was surgically obtained as part of a clinical trial (NTC03299946) for neoadjuvant cabozantinib and nivolumab previously described [38]. The surgical specimen was immediately embedded in OCT, frozen and a 10 μm cryosection was placed in a Visium Gene Expression slide (10x Genomics).

### 4.2 ST library preparation

Briefly, following tissue permeabilization optimization, according to 10x Genomics instructions, samples were fixed in methanol, stained (H&E) and imaged. Sequencing libraries were prepared using the Visium Spatial Gene Expression Reagent Kit (10x Genomics), following manufacturer’s instructions, and sequenced on an Illumina NovaSeq.

### 4.3 SpaceMarkers algorithm

Here we describe the SpaceMarkers algorithm to identify genes associated with nonlinear effects of latent feature interactions. To facilitate exposition, we will refer to the spatial component of the latent features as ”patterns”.

#### Modeling pattern interactions in the residual space

We assume a generic latent space representation model where the ST data matrix *D* is factorized into two low-rank matrices *A* and *P*. Consequently, the matrix product *AP* is a low-rank approximation of the high-dimensional spatial RNAseq data, accounting for all linear combinations of the latent patterns such that

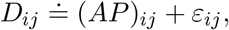

where measurement noise *ε_ij_* are independent and normally distributed with zero mean (see [24] for the CoGAPS-specific model). However, this model does not take into account any molecular changes resulting from inter-pattern interactions. We introduce a model where *D* is the sum of linear pattern effects represented by *AP* as well as unknown nonlinear effects due to inter pattern interactions represented by a matrix *f* (*A, P*) such that

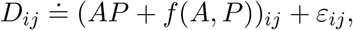

where the measurement noise *ε_ij_* are independent and normally distributed with zero mean and variance 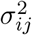. If we assume that the nonlinear effects *f* are known, we can modify the CoGAPS algorithm to account for the corresponding additional terms. However, since the nonlinear effects are unknown and may differ for each pair of patterns, this approach is infeasible. Our aim in this paper is not the exact characterization of the nonlinear effects, but to identify genes which exhibit higher nonlinear effects when two patterns are interacting. To this end, we use CoGAPS with the default settings and analyze the residual space of the CoGAPS factorization results. That is, we compare the CoGAPS residuals

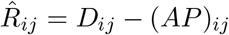

in regions where two patterns interact (i.e., have overlapping influence) versus regions where each pattern has exclusive influence. To identify the genes associated with the nonlinear interactions between a given pair of patterns, we first identify hotspots of pattern influence for each pattern. If both patterns have overlapping influence in a spot, they are deemed to be interacting in that spot. The CoGAPS residuals are computed in the interacting regions as well as in regions where each pattern is individually active. When the null hypothesis of non-interaction between the patterns is true, the residuals have no dependence on underlying regions (interacting or exclusive). On the other hand, genes associated with higher CoGAPS residuals in the interacting regions compared with the regions with exclusive pattern influence from either pattern show a strong dependence on spatial overlap between the patterns, and thus reject the null hypothesis. These genes constitute the SpaceMarkers, markers of spatial interaction between the two patterns in question. Focusing on strictly higher residuals avoids the confounding factors from decreased gene expression due to heterogeneous spot populations compared to homogeneous ones.

#### Identifying regions of pattern influence and pattern interaction

For each spatially resolved pattern, we identify its region of influence by way of an outlier detection method using a Gaussian kernel-based spatial smoothing. Given the pattern intensity *p*(*s_i_*) associated with a *i*-th spot *s_i_* = (*x_i_, y_i_*) in the sample, we calculate the spatially smoothed pattern intensities by using the leave-one-out method

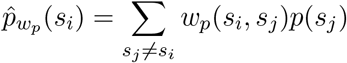

where the spatial kernel is a Gaussian kernel

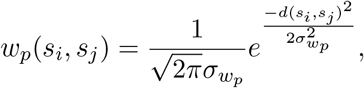

where 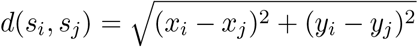 is the distance between the *i*-th and *j*-th spots, and *σ_w_p__* is the kernel width. We used the Smooth.ppp function in the R package spatstat [39] to perform the smoothing. We obtain a null-distribution by applying the kernel-based smoothing to spatially permuted pattern values (by pseudorandomly assigning spot locations (*n_perm_* = 100)). This null-distribution is assumed to be normal, and we obtain the sample mean 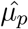 and standard deviation 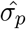 for each pattern. We identify the pattern’s region of influence as the set of spots with outliers

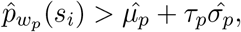

where *τ_p_* is the outlier threshold for the pattern. The optimal values of the kernel width *w_p_* and outlier threshold *τ_p_* are the arguments that minimize the spatial autocorrelation (Moran’s I) of the residuals

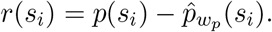

The optimal kernel width *w_p_* for each pattern is the value which minimizes the Moran’s I in the residuals over all spots in the sample. Subsequently, the optimal outlier threshold *τ_p_* minimizes spatial autocorrelation of the residuals *r*(*s_i_*) over the spots contained in the resulting region of pattern influence. If a spot is influenced by two or more patterns, these patterns are said to be interacting in such a spot. For each pattern pair of interest, the set of all such spots is defined as their interacting region.

#### Statistical test to identify genes associated with pattern interactions

For a given pair of patterns *p*_1_ and *p*_2_ with a substantial regions of exclusive pattern influence and pattern interaction, we define three subregions characterized by

- The spots with *p*_1_ influence and no *p*_2_ influence.
- The spots with *p*_2_ influence with no *p*_1_ influence.
- The spots with overlapping influence from both *p*_1_ and *p*_2_.

The elements from each row of 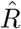 corresponding to the subregions described above denote the CoGAPS residuals in the respective subregions. For each gene (row) *i*, we perform a non-parametric Kruskal-Wallis test [40] for stochastic dominance of the CoGAPS residuals in at least one of the three subregions, with a posthoc Dunn’s test [41] to ascertain the relative dominance between the respective subregions. Of particular interest to us are the genes which have statistically significantly higher CoGAPS residuals (FDR¡0.05) in the interacting region relative to the other two subregions as well as genes which exhibit statistically significantly higher CoGAPS residuals exclusively in the interacting region compared to at least one of the two other subregions.

#### Code Availability

The SpaceMarkers package is available on Github at https://www.github.com/FertigLab/SpaceMarkers. All results in this manuscript were obtained using version 0.1.

### 4.4 Multi-resolution CoGAPS analysis

The ST genes by spot counts data for each sample was filtered to remove genes and spots with no or constant signal and then log2 normalized. The final matrix size of the input data matrix **D** are noted in the table below. The element *D*ij represents the expression of the *i*-th gene in the *j*-th spot. The CoGAPS (version 3.5.8)[33] algorithm was run using the filtered and normalized counts data as input. Additionally, default CoGAPS parameters were used except for nIterations = 50,000, sparseOptimization = TRUE, distributed = single-cell, and nSets = 4. CoGAPS factorization results in two lower-dimensional matrices: an amplitude matrix (**A**) containing gene weights and a pattern matrix (**P**) containing corresponding spot weights estimated for a pre-specified number of latent features (nPatterns). On each of the input datasets, the algorithm was tested for a range of nPatterns.

| Sample | # genes | # spots | numPatterns | # patterns learned |
| --- | --- | --- | --- | --- |
| PDAC metastatic lymph node | 18,418 | 1,351 | 8 | 10 |
| Invasive Breast Ductal Carcinoma | 24,228 | 4,898 | 5 | 5 |
| Invasive Breast Ductal Carcinoma | 24,228 | 4,898 | 20 | 16 |
| HCC | 20,423 | 3,006 | 15 | 9 |

The pattern weights for each spot were plotted over the tissue to show association between a pattern and a tissue region. In highRes Breast cancer analysis, genes were assigned to the pattern they were most strongly associated with using the patternMarker function in CoGAPS (version) in R (version). The genes for each pattern were submitted to the Molecular Signatures Database and searched within the BIOCARTA, KEGG, and HALLMARK pathways [27–29]. Pathways were considered significant if FDR < 0.05.

### 4.5 Scatterpie visualizations

We use the **A** and **P** matrices in the CoGAPS result to represent each Visium spot as a combination of overlapping latent patterns. To this end, we calculate the fractional gene expression *FSE_kj_* in pattern *k* at spot *j* as

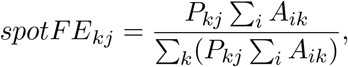

where *i* is the gene index. We use the ‘vizAllTopics‘ function from the ‘STdeconvolve‘ package [17] to visualize each spot as a pie chart showing the fractional gene expression in each pattern.

### 4.6 ProjectR analysis

The transfer learning method, ProjectR, was used to project the spatial patterns from the HCC sample onto matched scRNAseq data from the same patient. The R package projectR (version) was used to project the *A* matrix of the CoGAPS result into the target dataset. The CoGAPS result object and the counts data from the matched scRNAseq dataset were used as input where FULL = TRUE. Each individual cell in the scRNAseq dataset is associated with the pattern with the highest projection. We limit the pattern association to the dominant patterns in the spatial data, namely Patterns 1,2, and 8.

### 4.7 Gene Set Enrichment Analysis using MsigDB

For each gene list query corresponding to SpaceMarkers for pairs of patterns, we compute their overlaps with gene sets belonging to the HALLMARK, BIOCARTA and KEGG pathways in MsigDB [27–29], and report statistically significant overlaps (FDR¡0.05).

## Supporting information

High resolution version of Figure 1

High resolution version of Figure 2

High resolution version of Figure 3

High resolution version of Figure 4

High resolution version of Figure 5

High resolution version of Figure S1

High resolution version of Figure S2

High resolution version of Figure S3

High resolution version of Figure S4

## 5 Data availability

Spatial transcriptomics and scRNA-seq data are undergoing submission while the manuscript is under review.

## 6 Code availability

SpaceMarkers is implemented in the open-source R package SpaceMarkers, with source code freely available at https://www.github.com/FertigLab/SpaceMarkers. Additional code used for analysis in this paper is available at https://www.github.com/atuldeshpande/SpaceMarkers-paper. All results in this manuscript were obtained using SpaceMarkers version 0.1.

## 7 Acknowledgements

We thank Jennifer Durham for facilitating the sample acquisition and data archival process, and Ana Cordova for providing valuable inputs for improving the visualizations. This work was supported by an NCI F31CA268724-01 (to D.N.S), NIH K99 NS122085 from BRAIN Initiative in partnership with the National Institute of Neurological Disorders (to G.S.O); Kavli NDS Distinguished Postdoctoral Fellowship (to G.S.O), Johns Hopkins Provost Postdoctoral Fellowship (to G.S.O), R01CA138264 (to A.V.F.), R01DE027809 (to A.V.F.), U01CA212007 (to A.V.F.), R50CA243627 (to L.D.), NIH U01CA212007 (to A.S.P.), NIH R01CA138264 (to A.S.P.), NIH P01-CA247886-01A1 (to E.M.J.), SU2C/AACR DT-14-14 (to E.M.J.), Lustgarten Foundation grant (to E.M.J.), Lustgarten Foundation Pancreatic Cancer Research grant (to L.T.K.), The Sol Goldman Pancreatic Cancer Research Center grant (to L.T.K.), P01CA247886-01A1 (to L.T.K.)., the Emerson Cancer Research Fund (to E.M.J., E.J.F., W.J.H), an Allegheny Health Network (AHN) grant (to E.J.F.), U01CA212007 (to E.J.F.), U01CA253403 (to E.J.F.), the JHU Discovery Award (to E.J.F.), and SPORE GI P50CA062924-24A1 (to E.M.J, E.J.F., L.T.K, and L.Z.).

## Supplementary Figures

**Figure S1.**
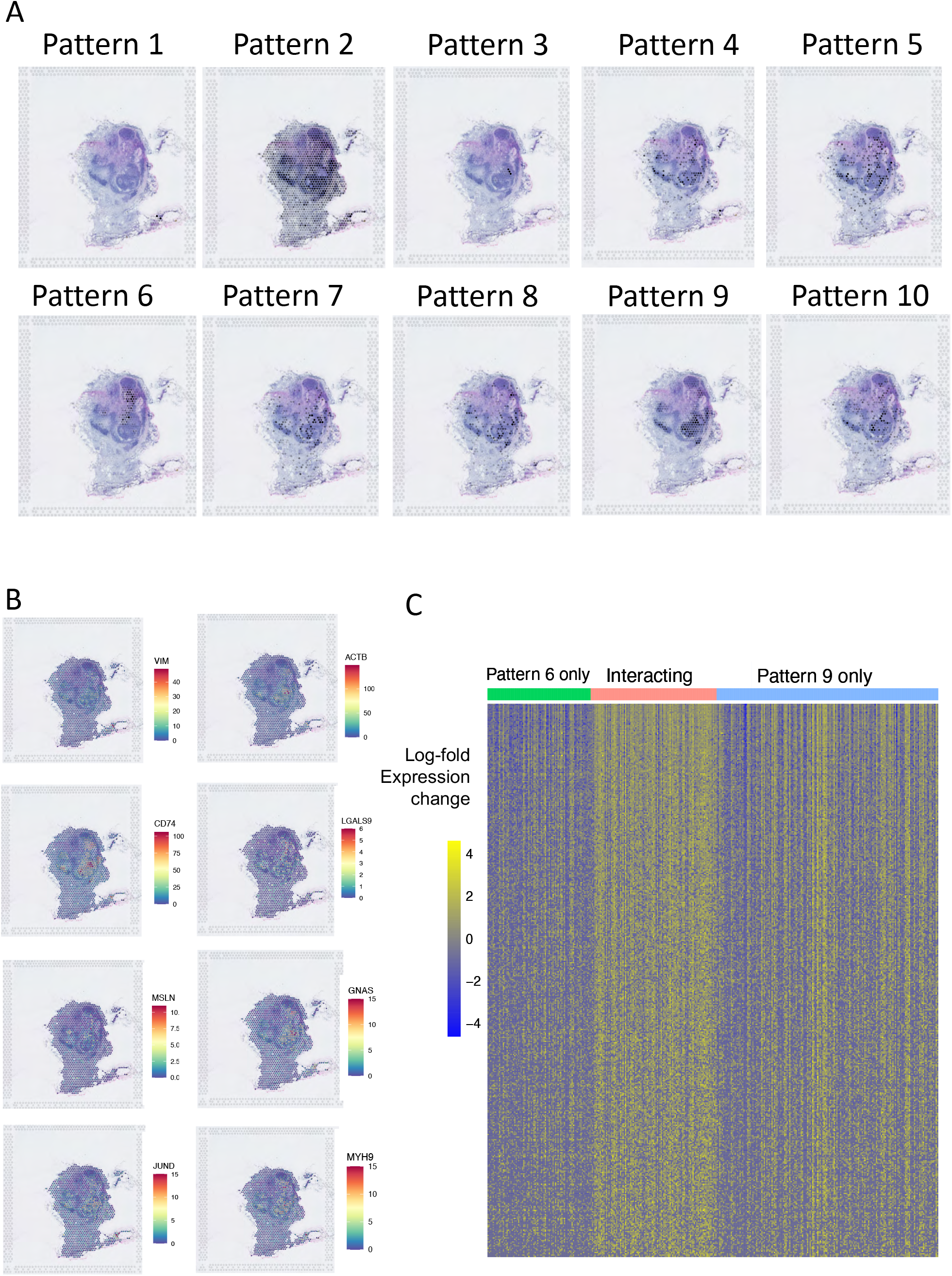
Analysis of lymph node tissue with metastatic PDAC (Related to Figure 2). **A.** Intensity maps of CoGAPS patterns identified in the lymph tissue. Pattern 6 shows high activity levels in the region annotated as metastatic PDAC and Pattern 9 shows high activity levels in the region associated with the surrounding immune cells. **B.** Spatial expression profile of the genes with expression box plots in Figure 2C. shows higher expression on the interface of the metastatic PDAC and immune cells. **C.** Expression heatmap of top 500 genes identified as SpaceMarkers in the regions with Pattern 6 and Pattern 9 interacting compared to regions with exclusive influence from Pattern 6 and Pattern 9 respectively. Complete gene list provided in Table S2.

**Figure S2.**
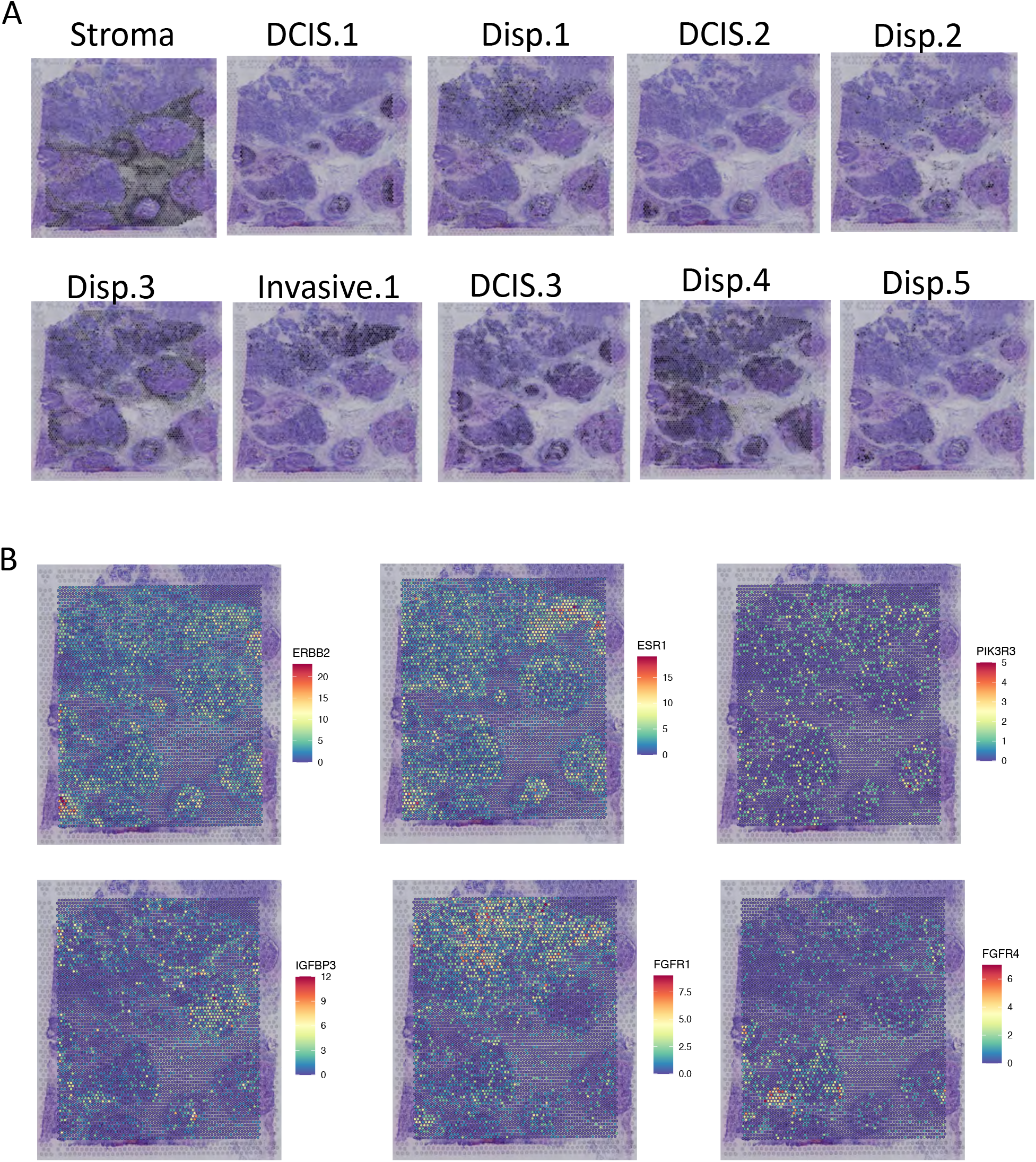
Analysis of high dimensional CoGAPS factorization of the breast cancer tissue (Related to Figure 4). **A.** Spatial intensity plots of CoGAPS patterns not shown in Figure 4. **B.** Spatial expression heatmap of select genes demonstrating the heterogeneity in the tissue sample. Whereas *ERBB2*, *ESR1* and *PIK3R3* are universally associated with all DCIS and Invasive lesions, *FGFR1* (Invasive.2), *IGFBP3* (DCIS.5) and *FGFR4* (DCIS.6) are associated with patterns associated with specific lesions.

**Figure S3.**
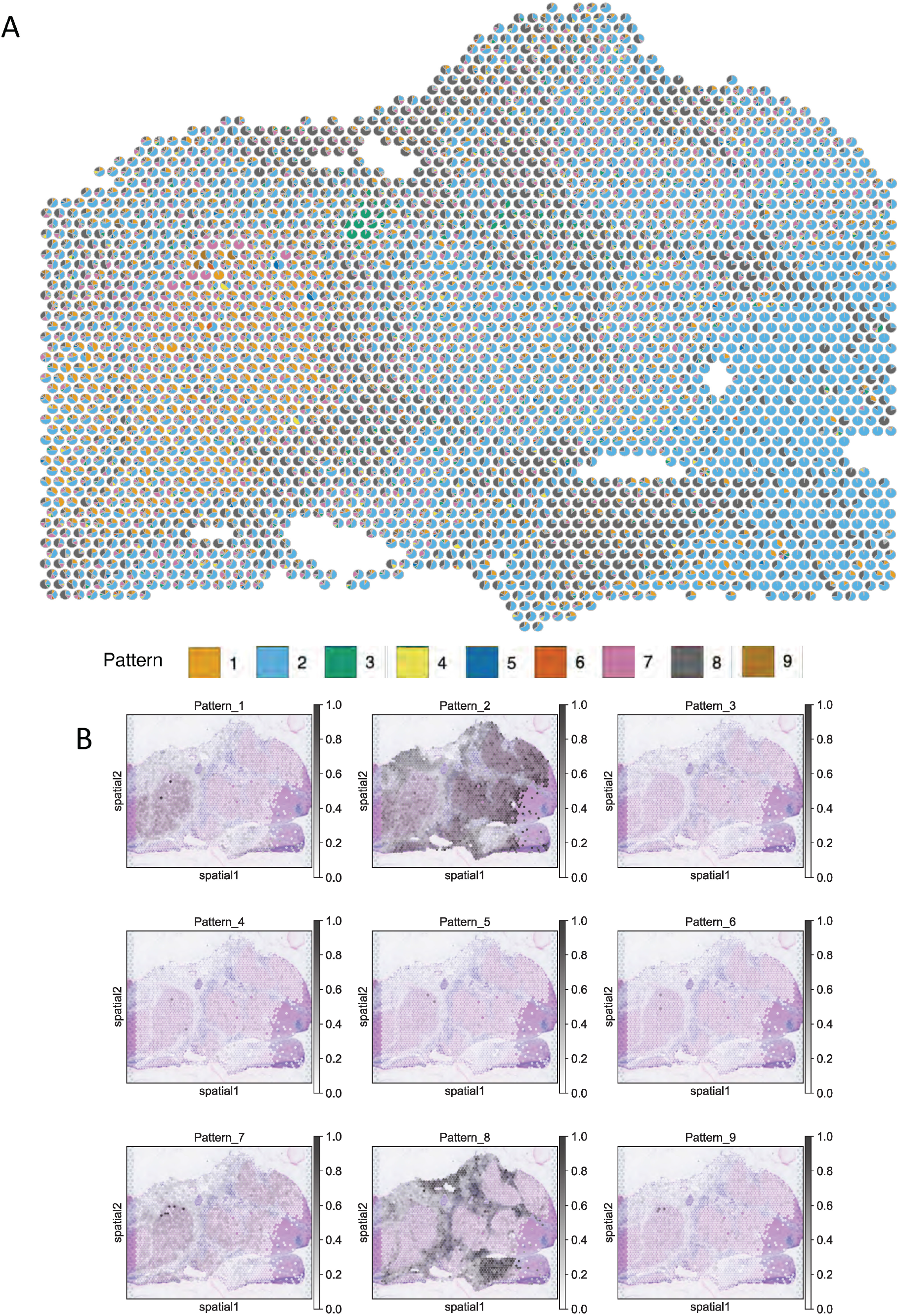
(CoGAPS learned spatial patterns from HCC tumor sample. Related to Figure 5). **A.** Scatterpie visualization showing the relative activity levels of all 9 patterns in each spot shows Patterns 1, 2, and 8 being the dominant patterns in the tissue. **B.** Spatial intensity plots of individual patterns normalized to sum to 1 in each spot also shows that Pattern 1, 2, and 8 explain most of the gene signature in the tissue.

**Figure S4.**
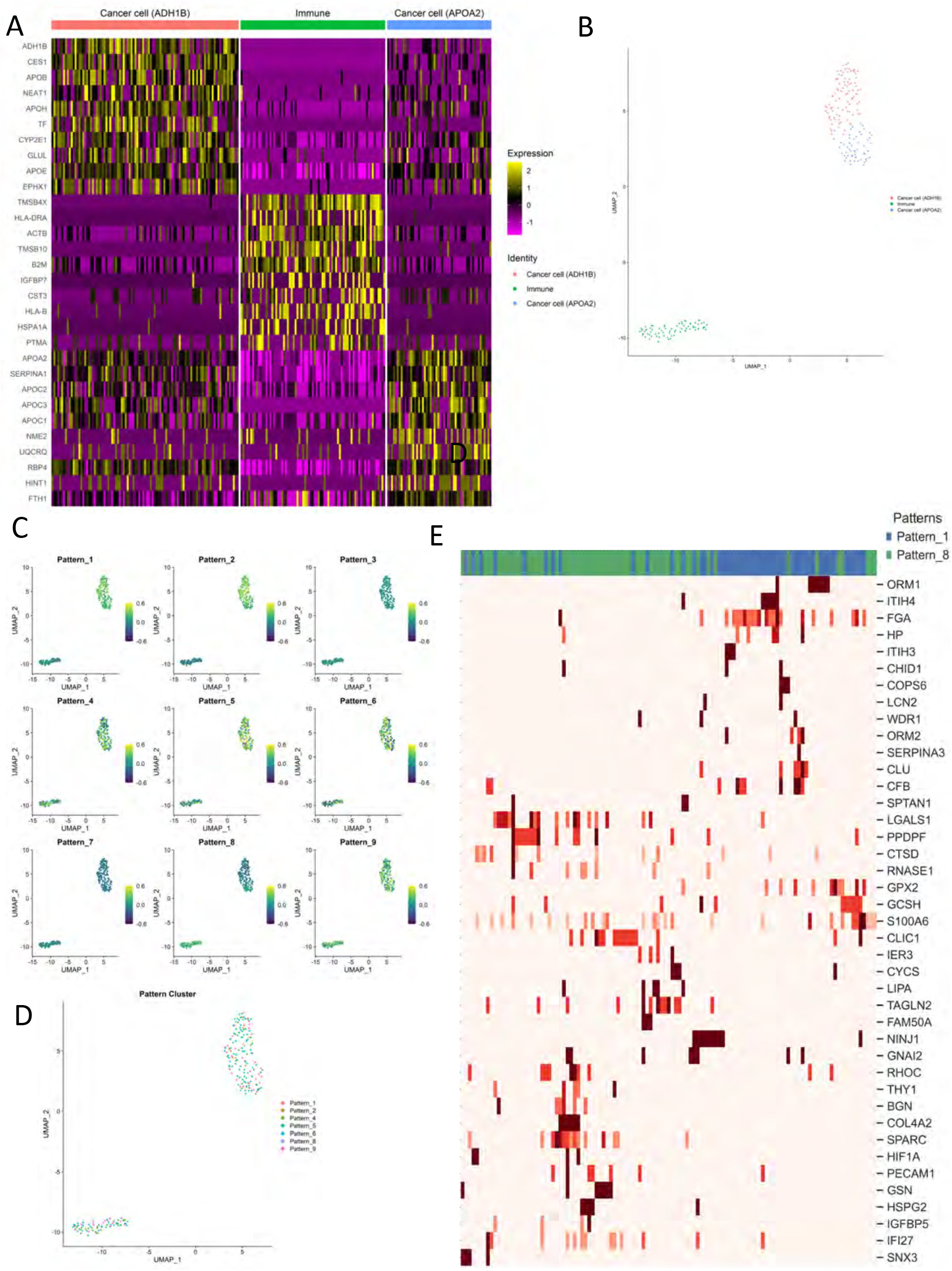
Analysis of SpaceMarkers in matched scRNAseq data (Related to Figure 5). **A.** Unsupervised clustering of the scRNAseq data, using Seurat, reveals two distinct tumor cell clusters and an immune cell cluster. Expression heatmap of select cancer and immune markers shows heterogeneity in the sample. **B.** UMAP plots showing projections of the spatial patterns learned in Figure S3 onto the scRNAseq data using ProjectR transfer learning. **C.** Each cell in the scRNAseq data is associated with the pattern with the highest projection in the cell as seen in panel B. However, since Patterns 1, 2, and 8 are dominant in the matched ST data, we assign cells to those three patterns as shown in Figure 5A. **D.** Expression heatmap of SpaceMarkers associated with interactions between Pattern 1 and Pattern 8 provide the spatial context of the individual cells.

## Notes

https://www.github.com/FertigLab/SpaceMarkers

https://www.github.com/atuldeshpande/SpaceMarkers-paper

### Competing Interest Statement

EJF is on the Scientific Advisory Board of Viosera Therapeutics and is a paid consultant for Merck and Mestag Therapeutics. WJH reports royalties from Rodeo/Amgen; grants from Sanofi and NeoTX; and consulting fees from Exelixis outside the submitted work. EMJ reports other support from Abmeta, personal fees from Genocea, personal fees from Achilles, personal fees from DragonFly, other support from Parker Institute, grants and other support from Lustgarten, personal fees from Carta, grants and other support from Genentech, grants and other support from AstroZeneca, personal fees from NextCure, and grants and other support from Break Through Cancer. SRW, CRU, JC, AH, and ZWB are equity stockholders and employees of 10x Genomics. LZ receives grant support from Bristol-Meyer Squibb, Merck, AstraZeneca, iTeos, Amgen, NovaRock, Inxmed, and Halozyme. LZ is a paid consultant/Advisory Board Member at Biosion, Alphamab, NovaRock, Ambrx, Akrevia/Xilio, QED, Natera, Novagenesis, Snow Lake Captials, Tempus, Amberston, and Mingruizhiyao. LZ holds shares at Alphamab and Mingruizhiyao. MY reports grants and research support from: Bristol-Myers Squibb, Incyte, Genentech, and honoraria from Genentech, Exelixis, Eisai, AstraZeneca, Replimune, Hepion. CSM reports research funding from Pfizer, Astrazeneca, BMS, GSK/Tesaro, and serves on the advisory boards of Bristol Myers Squibb (not paid), Seattle Genetics, Genomic Health, Athenex. RAA receives research support from Bristol Myers Squibb, RAPT therapeutics, Stand up to Cancer, and the National Institutes of Health and serves on the advisory boards for Bristol Myers Squibb, Merck SD, and AstraZeneca. ASP is a consultant to AsclepiX Therapeutics and CytomX Therapeutics; he is the founder and Chief Scientific Advisor of AsclepiX Therapeutics; he receives research grants from AstraZeneca and Boehringer Ingelheim. The terms of these arrangements are being managed by the Johns Hopkins University in accordance with its conflict-of-interest policies.

## References

1. H. R. Seo, “Roles of tumor microenvironment in hepatocelluar carcinoma,” Curr. Med. Chem., vol. 11, p. 82, June 2015.

2. E. F. Davis-Marcisak, A. Deshpande, G. L. Stein-O’Brien, W. J. Ho, D. Laheru, E. M. Jaffee, E. J. Fertig, and L. T. Kagohara, “From bench to bedside: single-cell analysis for cancer immunotherapy,” Cancer Cell, July 2021.

3. S. Juengpanich, W. Topatana, C. Lu, D. Staiculescu, S. Li, J. Cao, J. Lin, J. Hu, M. Chen, J. Chen, and X. Cai, “Role of cellular, molecular and tumor microenvironment in hepatocellular carcinoma: Possible targets and future directions in the regorafenib era,” Int. J. Cancer, vol. 147, pp. 1778–1792, Oct. 2020.

4. T. L. Whiteside, “The tumor microenvironment and its role in promoting tumor growth,” Oncogene, vol. 27, pp. 5904–5912, Oct. 2008.

5. R. Dhanasekaran, V. Baylot, M. Kim, S. Kuruvilla, D. I. Bellovin, N. Adeniji, A. Rajan Kd, I. Lai, M. Gabay, L. Tong, M. Krishnan, J. Park, T. Hu, M. A. Barbhuiya, A. J. Gentles, K. Kannan, P. T. Tran, and D. W. Felsher, “MYC and twist1 cooperate to drive metastasis by eliciting crosstalk between cancer and innate immunity,” Elife, vol. 9, Jan. 2020.

6. L.-Q. Fu, W.-L. Du, M.-H. Cai, J.-Y. Yao, Y.-Y. Zhao, and X.-Z. Mou, “The roles of tumor-associated macrophages in tumor angiogenesis and metastasis,” Cell. Immunol., vol. 353, p. 104119, July 2020.

7. C. Wei, C. Yang, S. Wang, D. Shi, C. Zhang, X. Lin, Q. Liu, R. Dou, and B. Xiong, “Crosstalk between cancer cells and tumor associated macrophages is required for mesenchymal circulating tumor cell-mediated colorectal cancer metastasis,” Mol. Cancer, vol. 18, p. 64, Mar. 2019.

8. B. Chaudhary and E. Elkord, “Regulatory T cells in the tumor microenvironment and cancer progression: Role and therapeutic targeting,” Vaccines (Basel), vol. 4, Aug. 2016.

9. T. Li, T. Liu, W. Zhu, S. Xie, Z. Zhao, B. Feng, H. Guo, and R. Yang, “Targeting MDSC for Immune-Checkpoint blockade in cancer immunotherapy: Current progress and new prospects,” Clin. Med. Insights Oncol., vol. 15, p. 11795549211035540, Jan. 2021.

10. A. E. Place, S. Jin Huh, and K. Polyak, “The microenvironment in breast cancer progression: biology and implications for treatment,” Breast Cancer Res., vol. 13, p. 227, Nov. 2011.

11. S. D. Soysal, A. Tzankov, and S. E. Muenst, “Role of the tumor microenvironment in breast cancer,” Pathobiology, vol. 82, pp. 142–152, Sept. 2015.

12. N. Rao, S. Clark, and O. Habern, “Bridging genomics and tissue pathology,” Genetic Engineering & Biotechnology News, vol. 40, pp. 50–51, Feb. 2020.

13. S. Z. Wu, G. Al-Eryani, D. L. Roden, S. Junankar, K. Harvey, A. Andersson, A. Thennavan, C. Wang, J. R. Torpy, N. Bartonicek, T. Wang, L. Larsson, D. Kaczorowski, N. I. Weisenfeld, C. R. Uytingco, J. G. Chew, Z. W. Bent, C.-L. Chan, V. Gnanasambandapillai, C.-A. Dutertre, L. Gluch, M. N. Hui, J. Beith, A. Parker, E. Robbins, D. Segara, C. Cooper, C. Mak, B. Chan, S. Warrier, F. Ginhoux, E. Millar, J. E. Powell, S. R. Williams, X. S. Liu, S. O’Toole, E. Lim, J. Lundeberg, C. M. Perou, and A. Swarbrick, “A single-cell and spatially resolved atlas of human breast cancers,” Nat. Genet., vol. 53, pp. 1334–1347, Sept. 2021.

14. A. L. Ji, A. J. Rubin, K. Thrane, S. Jiang, D. L. Reynolds, R. M. Meyers, M. G. Guo, B. M. George, A. Mollbrink, J. Bergenstråhle, L. Larsson, Y. Bai, B. Zhu, A. Bhaduri, J. M. Meyers, X. Rovira-Clavé, S. T. Hollmig, S. Z. Aasi, G. P. Nolan, J. Lundeberg, and P. A. Khavari, “Multimodal analysis of composition and spatial architecture in human squamous cell carcinoma,” Cell, vol. 182, pp. 497–514.e22, July 2020.

15. A. Andersson, L. Larsson, L. Stenbeck, F. Salmén, A. Ehinger, S. Z. Wu, G. Al-Eryani, D. Roden, A. Swarbrick, A. Borg, J. Frisén, C. Engblom, and J. Lundeberg, “Spatial deconvolution of HER2-positive breast cancer delineates tumor-associated cell type interactions,” Nat. Commun., vol. 12, p. 6012, Oct. 2021.

16. D. M. Cable, E. Murray, L. S. Zou, A. Goeva, E. Z. Macosko, F. Chen, and R. A. Irizarry, “Robust decomposition of cell type mixtures in spatial transcriptomics,” Nat. Biotechnol., Feb. 2021.

17. B. F. Miller, F. Huang, L. Atta, A. Sahoo, and J. Fan, “Reference-free cell-type deconvolution of multi-cellular pixel-resolution spatially resolved transcriptomics data,” bioRxiv, 2021.

18. E. Zhao, M. R. Stone, X. Ren, J. Guenthoer, K. S. Smythe, T. Pulliam, S. R. Williams, C. R. Uytingco, S. E. B. Taylor, P. Nghiem, J. H. Bielas, and R. Gottardo, “Spatial transcriptomics at subspot resolution with BayesSpace,” Nat. Biotechnol., vol. 39, pp. 1375–1384, Nov. 2021.

19. J. Hu, X. Li, K. Coleman, A. Schroeder, N. Ma, D. J. Irwin, E. B. Lee, R. T. Shinohara, and M. Li, “SpaGCN: Integrating gene expression, spatial location and histology to identify spatial domains and spatially variable genes by graph convolutional network,” Nat. Methods, vol. 18, pp. 1342–1351, Nov. 2021.

20. M. Elosua-Bayes, P. Nieto, E. Mereu, I. Gut, and H. Heyn, “SPOTlight: seeded NMF regression to deconvolute spatial transcriptomics spots with single-cell transcriptomes,” Nucleic Acids Res., vol. 49, p. e50, May 2021.

21. B. A. Luca, C. B. Steen, M. Matusiak, A. Azizi, S. Varma, C. Zhu, J. Przybyl, A. Espín-Pérez, M. Diehn, A. A. Alizadeh, M. van de Rijn, A. J. Gentles, and A. M. Newman, “Atlas of clinically distinct cell states and ecosystems across human solid tumors,” Cell, vol. 184, no. 21, pp. 5482–5496.e28, 2021.

22. E. F. Davis-Marcisak, A. A. Fitzgerald, M. D. Kessler, L. Danilova, E. M. Jaffee, N. Zaidi, L. M. Weiner, and E. J. Fertig, “Transfer learning between preclinical models and human tumors identifies a conserved NK cell activation signature in anti-CTLA-4 responsive tumors,” Genome Med., vol. 13, p. 129, Aug. 2021.

23. G. L. Stein-O’Brien, B. S. Clark, T. Sherman, C. Zibetti, Q. Hu, R. Sealfon, S. Liu, J. Qian, C. Colantuoni, S. Blackshaw, L. A. Goff, and E. J. Fertig, “Decomposing cell identity for transfer learning across cellular measurements, platforms, tissues, and species,” Cell Systems, vol. 8, pp. 395–411.e8, May 2019.

24. E. J. Fertig, J. Ding, A. V. Favorov, G. Parmigiani, and M. F. Ochs, “CoGAPS: an R/C++ package to identify patterns and biological process activity in transcriptomic data,” Bioinformatics, vol. 26, pp. 2792–2793, Nov. 2010.

25. G. L. Stein-O’Brien, R. Arora, A. C. Culhane, A. V. Favorov, L. X. Garmire, C. S. Greene, L. A. Goff, Y. Li, A. Ngom, M. F. Ochs, Y. Xu, and E. J. Fertig, “Enter the matrix: Factorization uncovers knowledge from omics,” Trends Genet., vol. 34, pp. 790–805, Oct. 2018.

26. G. Sharma, C. Colantuoni, L. A. Goff, E. J. Fertig, and G. Stein-O’Brien, “projectR: an R/Bioconductor package for transfer learning via PCA, NMF, correlation and clustering,” Bioinformatics, vol. 36, pp. 3592–3593, June 2020.

27. A. Liberzon, A. Subramanian, R. Pinchback, H. Thorvaldsdóttir, P. Tamayo, and J. P. Mesirov, “Molecular signatures database (MSigDB) 3.0,” Bioinformatics, vol. 27, pp. 1739–1740, June 2011.

28. A. Subramanian, P. Tamayo, V. K. Mootha, S. Mukherjee, B. L. Ebert, M. A. Gillette, A. Paulovich, S. L. Pomeroy, T. R. Golub, E. S. Lander, and J. P. Mesirov, “Gene set enrichment analysis: a knowledge-based approach for interpreting genome-wide expression profiles,” Proc. Natl. Acad. Sci. U. S. A., vol. 102, pp. 15545–15550, Oct. 2005.

29. A. Liberzon, C. Birger, H. Thorvaldsdóttir, M. Ghandi, J. P. Mesirov, and P. Tamayo, “The Molecular Signatures Database (MSigDB) hallmark gene set collection,” Cell Syst, vol. 1, pp. 417–425, Dec. 2015.

30. S. Mohammadi, J. Davila-Velderrain, and M. Kellis, “A multiresolution framework to characterize single-cell state landscapes,” Nat. Commun., vol. 11, p. 5399, Oct. 2020.

31. T. Biancalani, G. Scalia, L. Buffoni, R. Avasthi, Z. Lu, A. Sanger, N. Tokcan, C. R. Vanderburg, Å. Segerstolpe, M. Zhang, I. Avraham-Davidi, S. Vickovic, M. Nitzan, S. Ma, A. Subramanian, M. Lipinski, J. Buenrostro, N. B. Brown, D. Fanelli, X. Zhuang, E. Z. Macosko, and A. Regev, “Deep learning and alignment of spatially resolved single-cell transcriptomes with Tangram,” Nat. Methods, vol. 18, pp. 1352–1362, Nov. 2021.

32. D. T. Pham, X. Tan, J. Xu, L. F. Grice, P. Y. Lam, A. Raghubar, and others, “stLearn: integrating spatial location, tissue morphology and gene expression to find cell types, cell-cell interactions and spatial trajectories within undissociated tissues,” bioRxiv, 2020.

33. T. D. Sherman, T. Gao, and E. J. Fertig, “CoGAPS 3: Bayesian non-negative matrix factorization for single-cell analysis with asynchronous updates and sparse data structures,” BMC Bioinformatics, vol. 21, p. 453, Oct. 2020.

34. L. Monteran and N. Erez, “The Dark Side of Fibroblasts: Cancer-Associated Fibroblasts as Mediators of Immunosuppression in the Tumor Microenvironment,” Front. Immunol., vol. 10, p. 1835, Aug. 2019.

35. M. Efremova, M. Vento-Tormo, S. A. Teichmann, and others, “CellPhoneDB: inferring cell–cell communication from combined expression of multi-subunit ligand–receptor complexes,” Nat. Protoc., 2020.

36. A. Giladi, M. Cohen, C. Medaglia, Y. Baran, B. Li, M. Zada, P. Bost, R. Blecher-Gonen, T.-M. Salame, J. U. Mayer, E. David, F. Ronchese, A. Tanay, and I. Amit, “Dissecting cellular crosstalk by sequencing physically interacting cells,” Nat. Biotechnol., vol. 38, pp. 629–637, May 2020.

37. D. Li, J. Ding, and Z. Bar-Joseph, “Identifying signaling genes in spatial single cell expression data,” Bioinformatics, Sept. 2020.

38. W. J. Ho, Q. Zhu, J. Durham, A. Popovic, S. Xavier, J. Leatherman, A. Mohan, G. Mo, S. Zhang, N. Gross, S. Charmsaz, D. Lin, D. Quong, B. Wilt, I. R. Kamel, M. Weiss, B. Philosophe, R. Burkhart, W. R. Burns, C. Shubert, A. Ejaz, J. He, A. Deshpande, L. Danilova, G. Stein-O’Brien, E. A. Sugar, D. A. Laheru, R. A. Anders, E. J. Fertig, E. M. Jaffee, and M. Yarchoan, “Neoadjuvant cabozantinib and nivolumab convert locally advanced hepatocellular carcinoma into resectable disease with enhanced antitumor immunity,” Nature Cancer, vol. 2, pp. 891–903, July 2021.

39. A. Baddeley, E. Rubak, and R. Turner, Spatial Point Patterns: Methodology and Applications with R. London: Chapman and Hall/CRC Press, 2015.

40. W. H. Kruskal and W. A. Wallis, “Use of ranks in One-Criterion variance analysis,” J. Am. Stat. Assoc., vol. 47, pp. 583–621, Dec. 1952.

41. O. J. Dunn, “Multiple comparisons using rank sums,” Technometrics, vol. 6, pp. 241–252, Aug. 1964.

