## Supplementary figures and images for "Uncovering the spatial landscape of molecular interactions within the tumor microenvironment through latent spaces"

### High resolution version of Figure 1

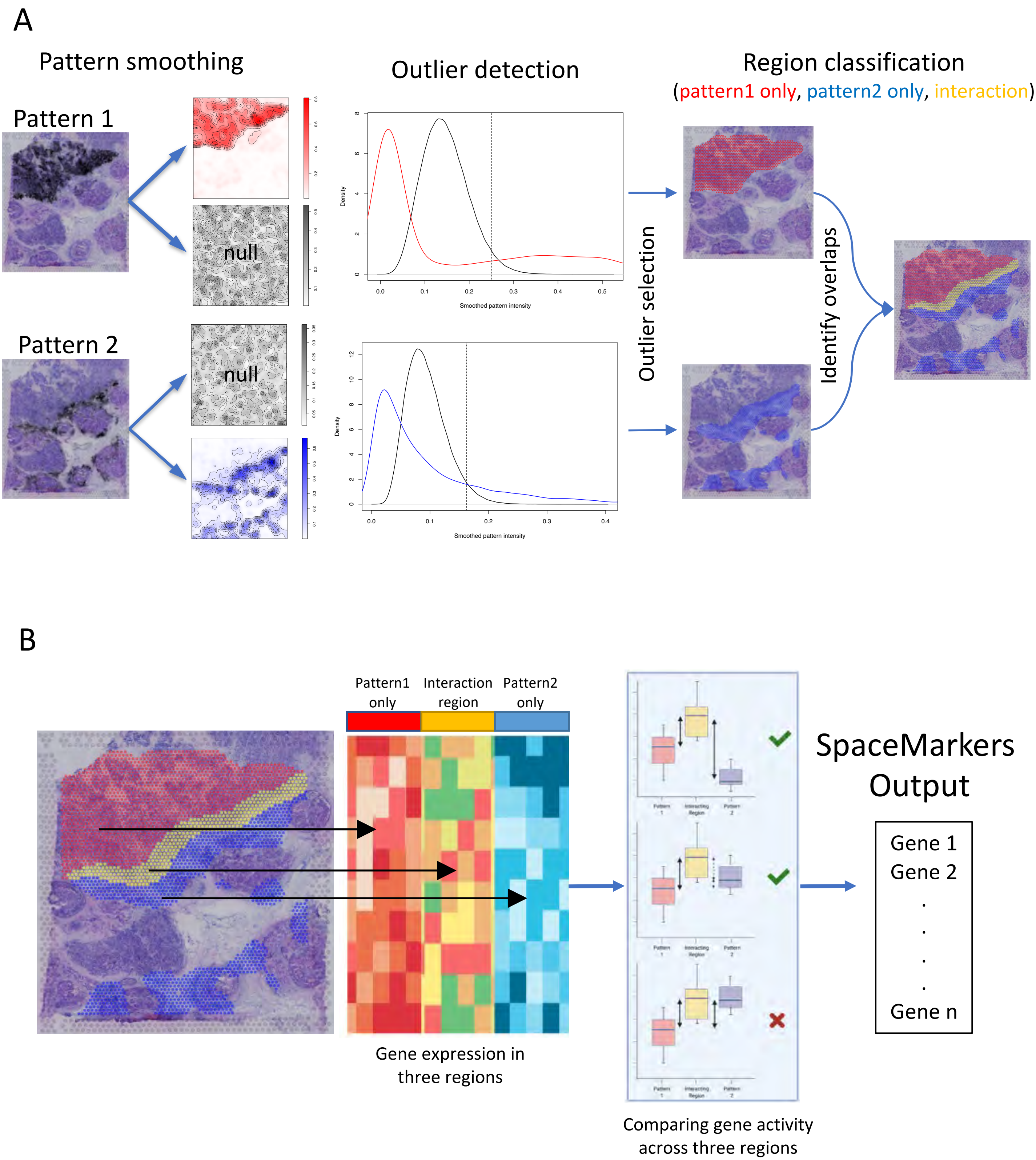

### High resolution version of Figure 3

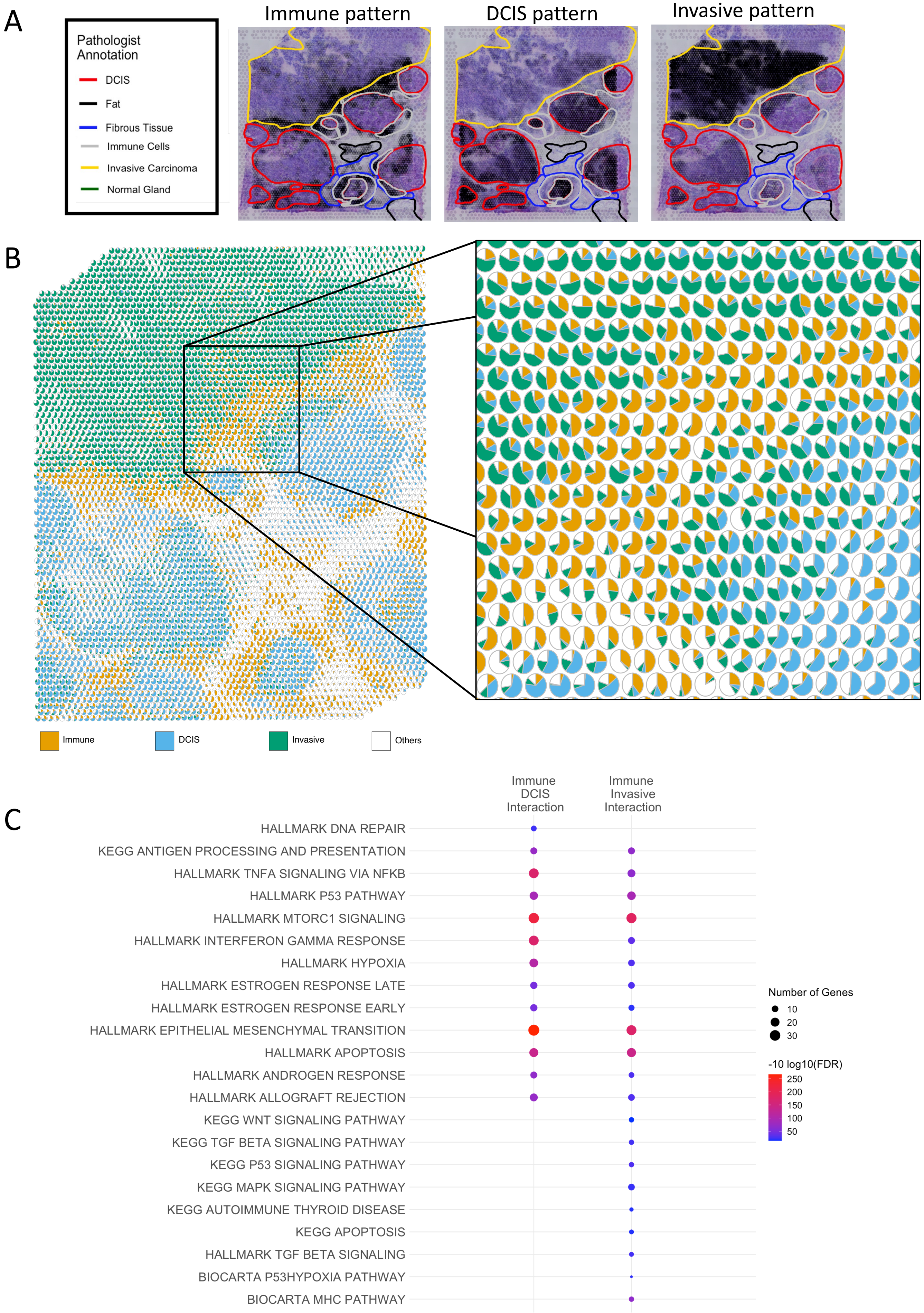

### High resolution version of Figure 4

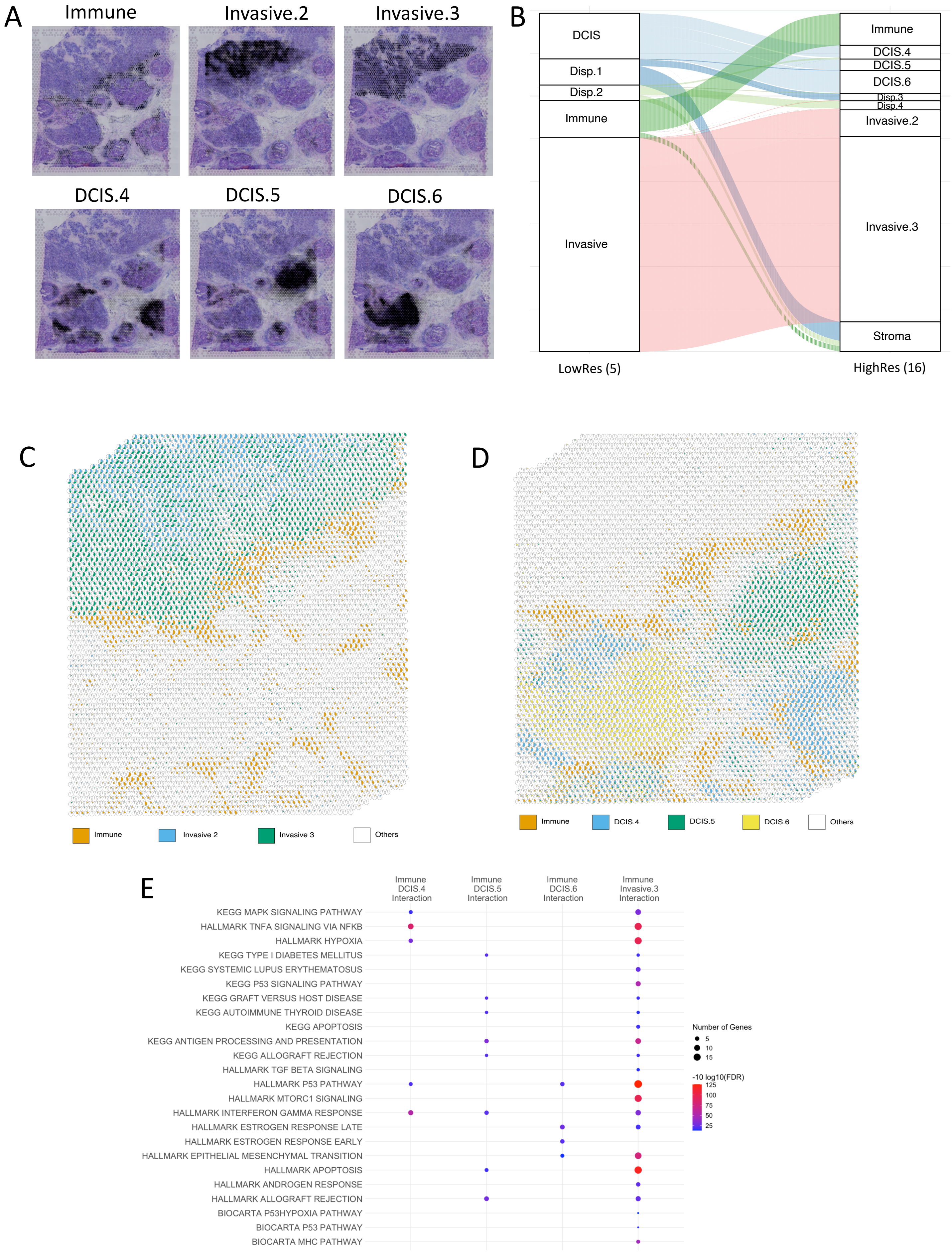

### High resolution version of Figure 5

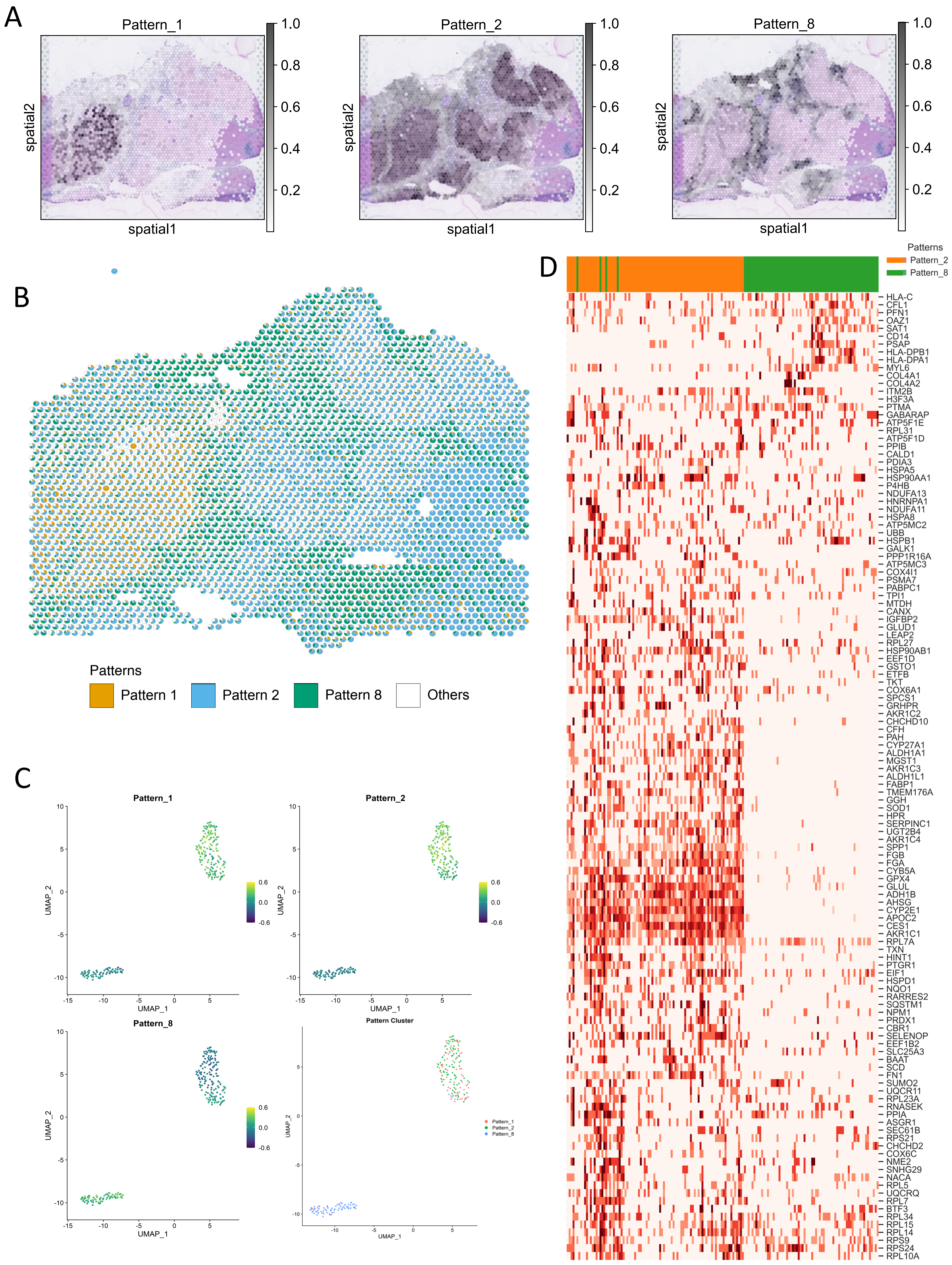

### High resolution version of Figure S1

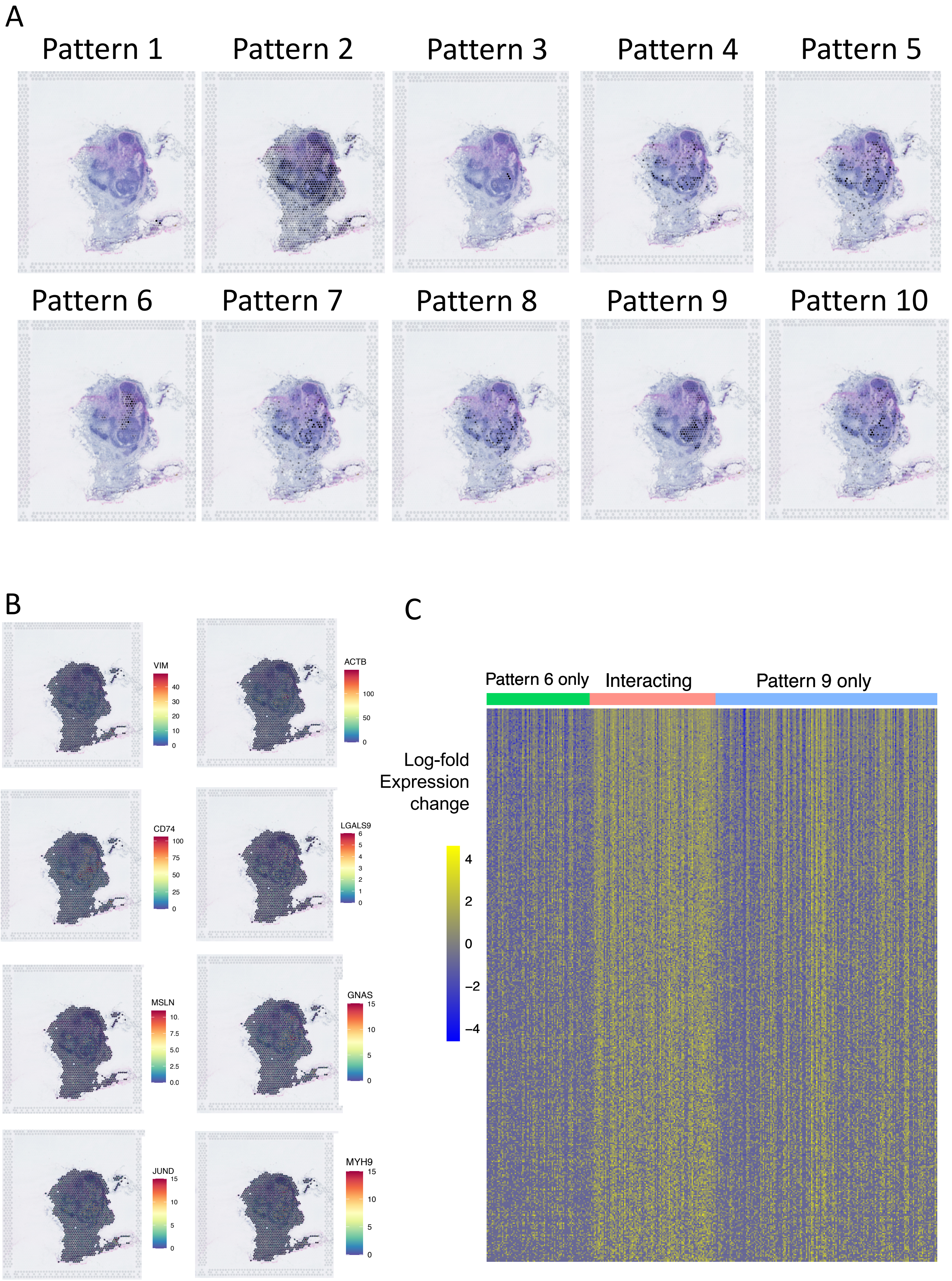

### High resolution version of Figure S2

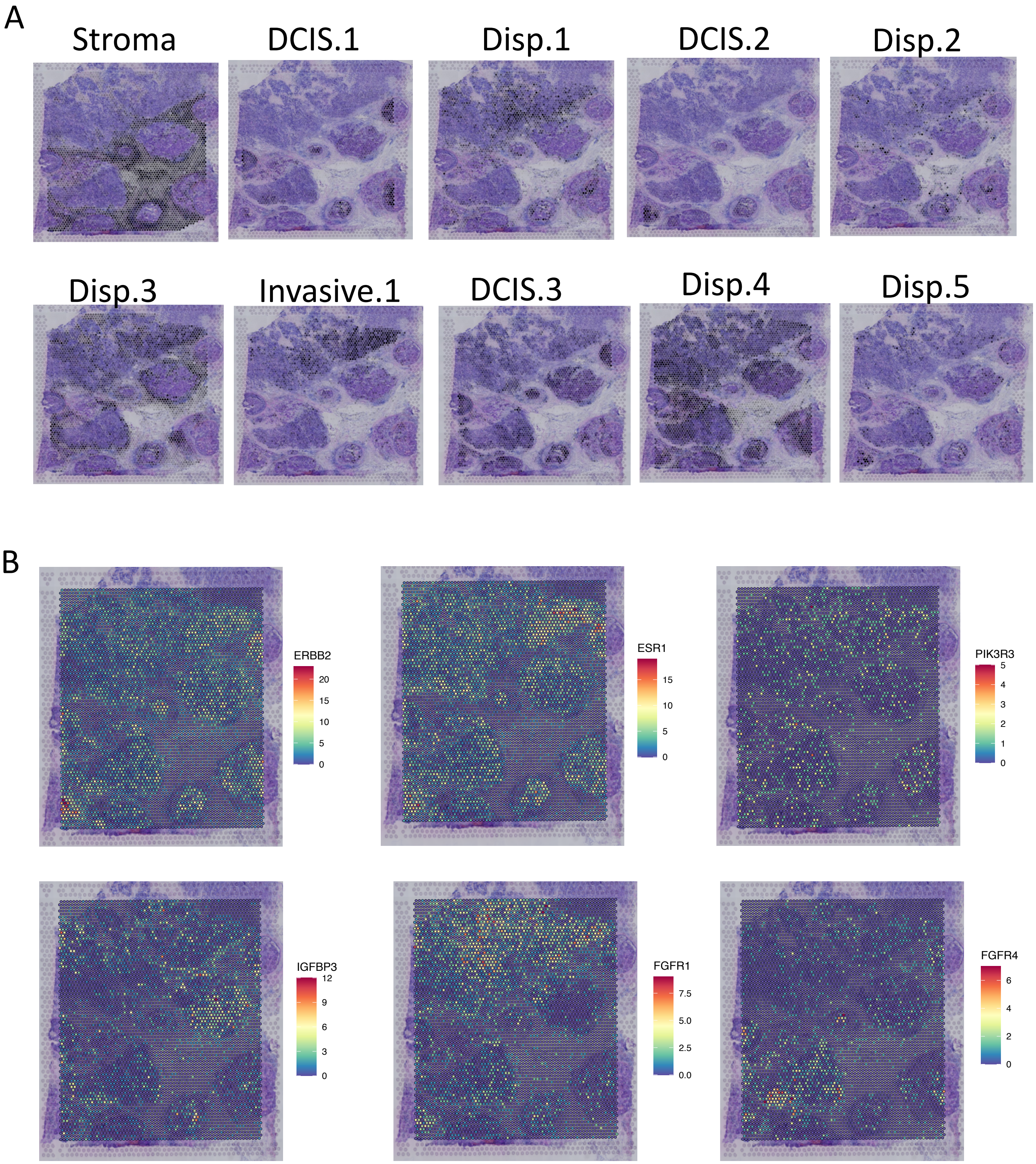

### High resolution version of Figure S3

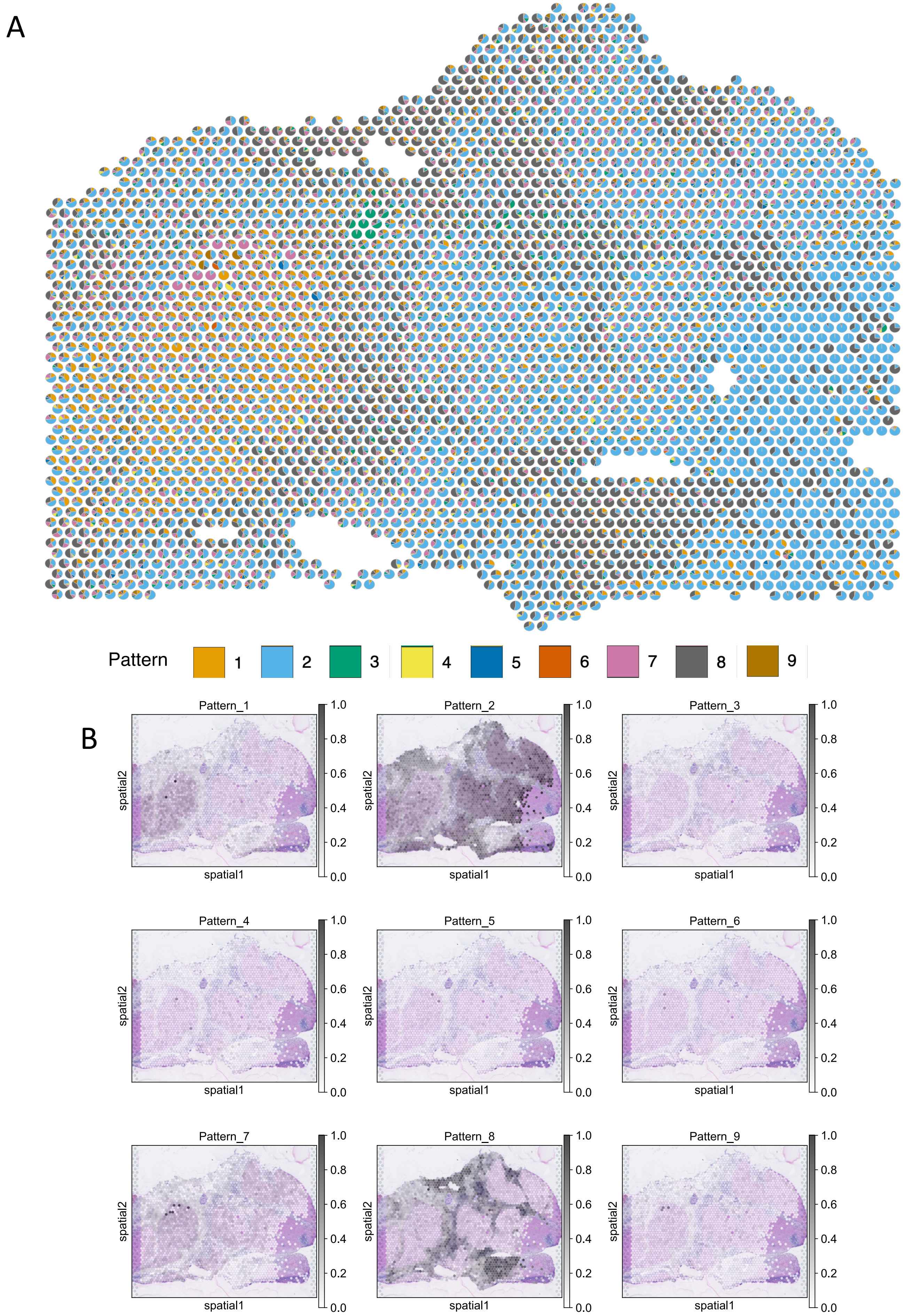

### High resolution version of Figure S4

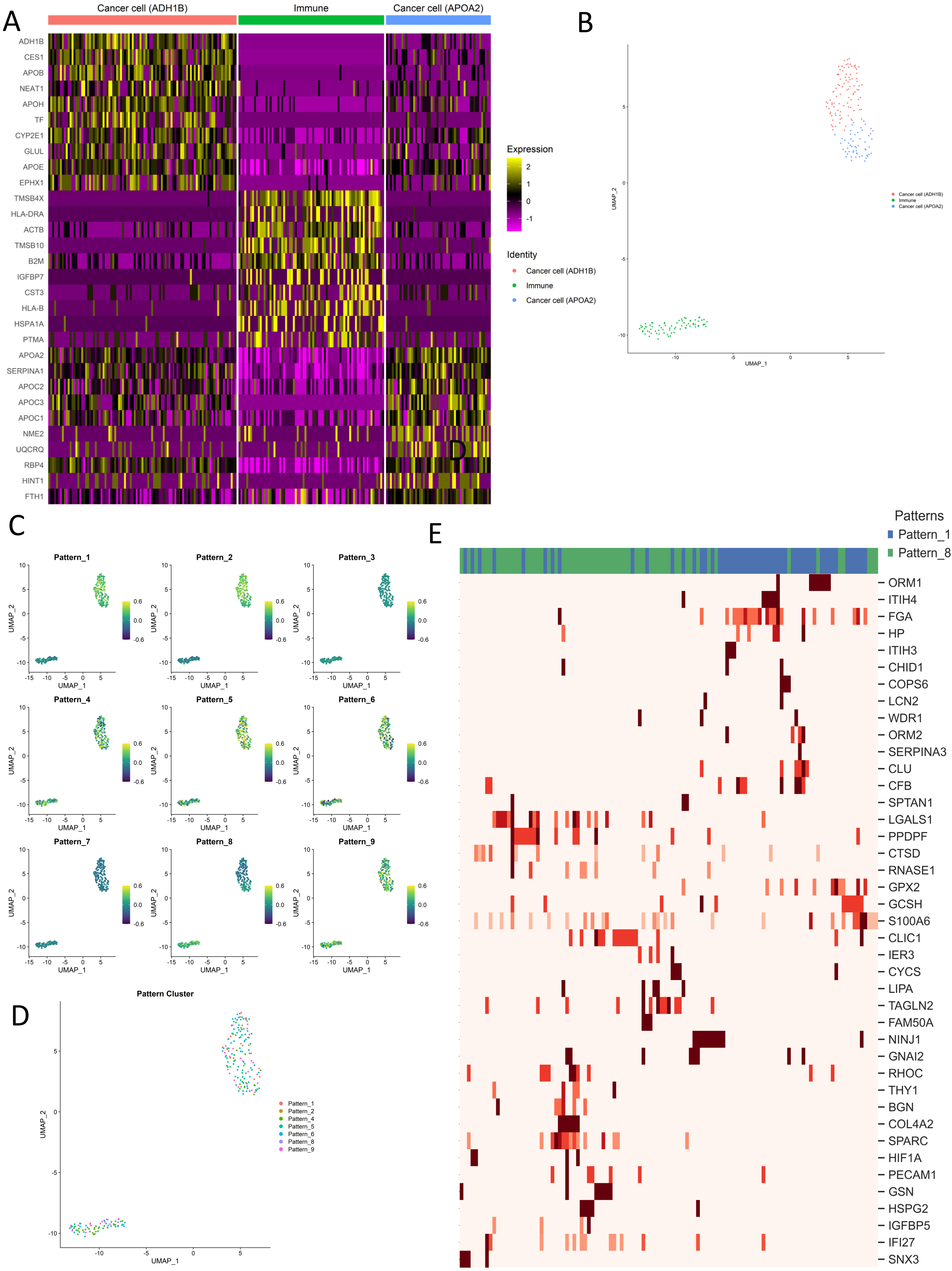
