## Supplementary material for "Uncovering the spatial landscape of molecular interactions within the tumor microenvironment through latent spaces": High resolution version of Figure 2

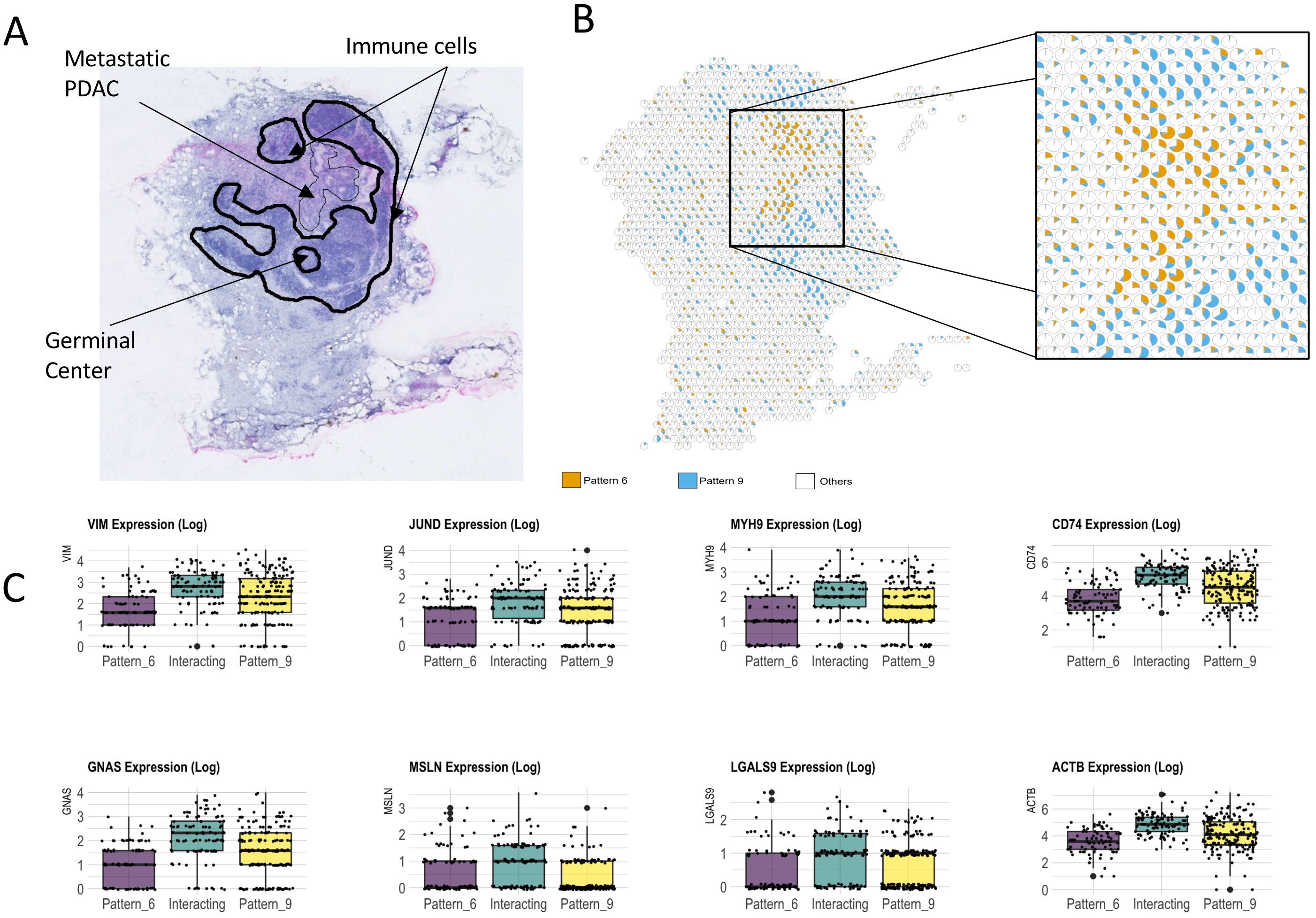


D

| Gene Set Name | # Genes in Gene Set | # Genes in Overlap | p-value | FDR q-value |
|---|---|---|---|---|
| HALLMARK MYC TARGETS V1 | 200 | 39 | 5.34E-35 | 1.41E-32 |
| HALLMARK ALLOGRAFT REJECTION | 200 | 29 | 1.70E-22 | 1.80E-20 |
| HALLMARK OXIDATIVE PHOSPHORYLATION | 200 | 24 | 6.73E-17 | 3.55E-15 |
| HALLMARK INTERFERON GAMMA RESPONSE | 200 | 21 | 8.69E-14 | 2.70E-12 |
| HALLMARK INTERFERON ALPHA RESPONSE | 97 | 13 | 2.21E-10 | 4.87E-09 |
| HALLMARK APICAL JUNCTION | 200 | 16 | 4.40E-09 | 8.01E-08 |
| HALLMARK MTORC1 SIGNALING | 200 | 16 | 4.40E-09 | 8.01E-08 |
| HALLMARK P53 PATHWAY | 200 | 16 | 4.40E-09 | 8.01E-08 |
| HALLMARK PI3K AKT MTOR SIGNALING | 105 | 12 | 7.03E-09 | 1.16E-07 |
| HALLMARK G2M CHECKPOINT | 200 | 15 | 3.18E-08 | 4.94E-07 |
| HALLMARK COMPLEMENT | 200 | 14 | 2.14E-07 | 3.14E-06 |
| HALLMARK APOPTOSIS | 161 | 12 | 8.08E-07 | 1.09E-05 |
| HALLMARK EPITHELIAL MESENCHYMAL TRANSITION | 200 | 12 | 7.72E-06 | 9.26E-05 |
| HALLMARK COAGULATION | 138 | 10 | 8.55E-06 | 1.00E-04 |
| HALLMARK UNFOLDED PROTEIN RESPONSE | 113 | 9 | 1.13E-05 | 1.29E-04 |
| HALLMARK INFLAMMATORY RESPONSE | 200 | 11 | 4.09E-05 | 4.32E-04 |
| HALLMARK GLYCOLYSIS | 200 | 10 | 1.98E-04 | 1.94E-03 |
| HALLMARK HYPOXIA | 200 | 10 | 1.98E-04 | 1.94E-03 |
| HALLMARK UV RESPONSE UP | 158 | 8 | 7.82E-04 | 6.07E-03 |
| HALLMARK MITOTIC SPINDLE | 199 | 9 | 8.41E-04 | 6.30E-03 |
| HALLMARK E2F TARGETS | 200 | 9 | 8.71E-04 | 6.30E-03 |
| HALLMARK MYOGENESIS | 200 | 9 | 8.71E-04 | 6.30E-03 |
| HALLMARK TNFA SIGNALING VIA NFKB | 200 | 9 | 8.71E-04 | 6.30E-03 |
| HALLMARK DNA REPAIR | 150 | 7 | 2.61E-03 | 1.66E-02 |
| HALLMARK IL2 STAT5 SIGNALING | 199 | 8 | 3.34E-03 | 2.00E-02 |
| HALLMARK ADIPOGENESIS | 200 | 8 | 3.44E-03 | 2.04E-02 |
| HALLMARK MYC TARGETS V2 | 58 | 4 | 5.62E-03 | 2.97E-02 |
